# Dissecting the molecular basis underlying mycobacterial cell-wall hydrolysis by the catalytic domains of *D29*LysA and *DS6A*LysA phage endolysins

**DOI:** 10.1101/2025.01.22.634383

**Authors:** Fernando Ceballos-Zúñiga, Laura Gálvez-Larrosa, Inés G. Muñoz, Lourdes Infantes, Inmaculada Pérez-Dorado

## Abstract

Mycobacteria encompass a broad range of microorganisms that cause infections with a significant impact on human health, resulting in millions of deaths each year. From tuberculosis and leprosy, caused by *Mycobacterium tuberculosis* and *Mycobacterium leprae*, respectively, to infections caused by emerging/opportunistic pathogens such as *Mycobacterium abscessus*. The battle to combat this health burden is further challenged by limitations in the treatments currently available and the rise of antimicrobial resistance. This underscores the need for new therapeutic strategies to combat these infections. Mycobacteriophage LysA endolysins are complex, multi-domain peptidoglycan hydrolases with reported antimicrobial relevance and the potential to treat mycobacterial infections. However, despite the therapeutic prospects of LysAs, our understanding of their mechanism of action remains limited. This study provides a comprehensive structural-functional analysis of the catalytic domains of two LysA endolysins encoded by the bacteriophages *D29* and *DS6A*, which are known to infect pathogenic mycobacteria, including *M. tuberculosis*. As part of this work, we have characterized the four catalytic domains present in both endolysins (*D29*N4/*D29*GH19 and *DS6A*GH19/*DS6A*Ami2B) both alone and in complex with PG analogues. To achieve this, we combined protein engineering, X-ray crystallography, small-angle X-ray scattering, and *in silico* tools. To our knowledge, this has led to the first experimental structures reported for mycobacteriophage endolysins, which reveals key aspects of peptidoglycan binding and hydrolysis by *D29*LysA and *DS6A*LysA lysins, as well as other homologous LysAs, including the hydrolase domains similar to those examined here. Altogether, this represents a significant step forward in understanding how mycobacterial cell-wall hydrolysis occurs by this important class of endolysins and opens the door to their future use in therapeutic applications as enzybiotics. Information that will allow the rational design of *a la carte* enzymes with optimized lytic properties against mycobacterial pathogens.

## 1. Introduction

The *Mycobacterium* genus contains over 170 recognized species, many of which cause infectious diseases (IDs) in humans, such as Buruli ulcer, leprosy, and tuberculosis (TB), caused by *Mycobacterium ulcerans* (*Mlc*), *Mycobacterium leprae*, and *Mycobacterium tuberculosis* (*Mtb*), respectively (Batt *et al*, 2020). Among them, human TB is the world’s most devastating human ID, being the leading cause of death globally caused by bacteria, according to the World Health Organization (Global Tuberculosis Report 2023, 2023). Buruli ulcer and leprosy are IDs associated with permanent deformation, disability, and stigma, causing over 200,000 new cases every year across more than 120 countries world (Global Leprosy (Hansen’s disease) Strategy 2021-2030 | 2, 2021). In addition, various non-tuberculous mycobacteria can cause IDs in immunocompromised individuals, such as the emerging pathogen *Mycobacterium abscessus* (Johansen *et al*, 2020). Great part of the success of mycobacteria as pathogens relies on their mycobacterial cell envelope (MCE), which consists of a plasma membrane surrounded by a complex structure formed by three distinct, covalently linked macromolecules: the outer membrane (OM) rich in mycolic acids (MA), arabinogalactan (AG), and peptidoglycan (PG) (**Fig. 1**) (Catalão *et al*, 2019). Altogether, this forms a very impermeable barrier essential for cell integrity that protects the bacilli from harsh conditions, making it one of the main reasons for antimicrobial resistance in mycobacteria (Catalão *et al*, 2019; Batt *et al*, 2020).

**Figure 1.**
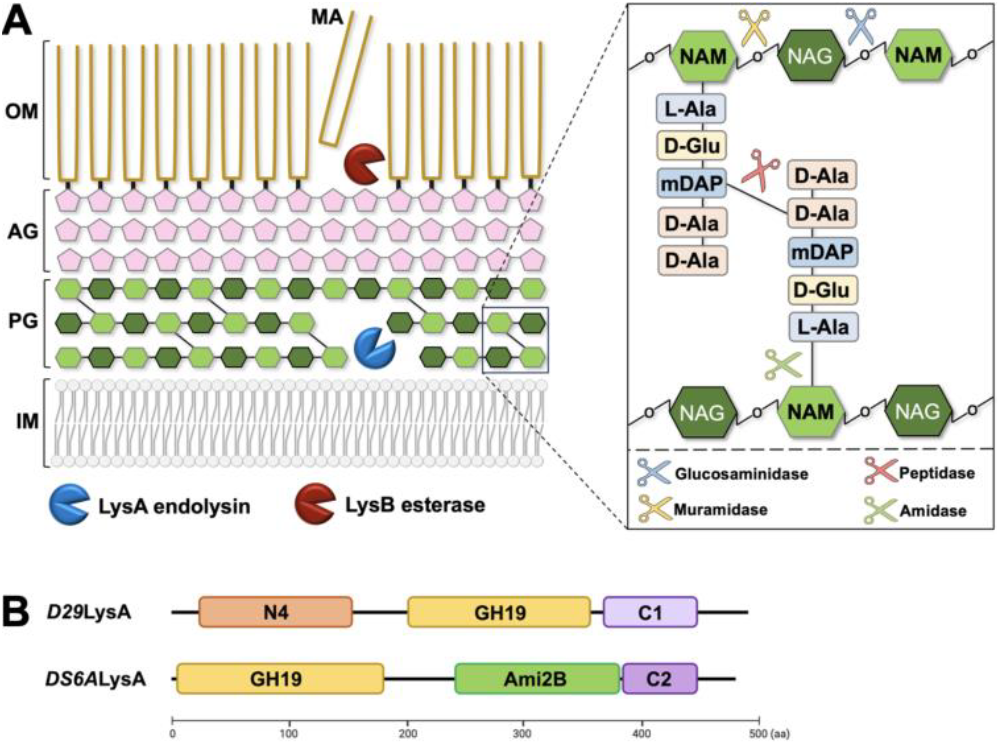
(A) Scheme representing the mycobacterial cell-wall architecture consisting of the inner membrane (IM) surrounded by the mycolyl-arabinogalactan-peptidoglycan (MA-AG-PG) complex (left). Mycobacteriophage LysA and LysB lysins targeting the PG and MA moieties are represented as blue and red packmen, respectively. Details of the peptidoglycan composition are shown (right), which consists of glycan strands formed by alternating N-acetylglucosamine (NAG) and N-acetylmuramic acid (NAM) and covalently cross-linked via peptide stems pending from NAM residues. The PG-hydrolase activities predicted in LysA endolysins are depicted as scissors. (B) Representation of the three-domain architecture predicted for *D29*LysA and *DS6*ALysA endolysins.

Among the MCE components, PG provides the mechanical strength to resist turgor pressure, while also playing a central role in regulating cellular shape, growth, and division (Batt *et al*, 2020; Vollmer *et al*, 2008). The PG molecule is built from linear saccharide strands of alternating units of *N*-acetylglucosamine (NAG) and *N*-acetylmuramic acid (NAM), connected by β-1,4 bonds (**Fig. 1**). These glycan strands are, in turn, covalently cross-linked via peptide stems formed by L-alanine (L-Ala), D-iso-glutamate (D-iso-Glu), meso-diaminopimelate (mDAP) acid, and D-alanine (D-Ala) present at NAM residues. Mycobacterial species exhibit characteristic modifications in both the glycan and peptide moieties, such as amidation of free carboxylic groups or glycolylation of NAM (*N*-glycolylmuramic acid or MurGlyc) (Catalão *et al*, 2019; Batt *et al*, 2020). While the former is important for PG biosynthesis, glycolylation may confer additional strength to the PG molecule, as well as resistance to lysozyme, thereby protecting mycobacteria against the innate immune system (Raymond *et al*, 2005). Moreover, mycobacteria exhibit a higher degree of peptide cross-linking (up to 80%) compared to other bacteria, resulting in a sturdier PG structure structure (Matsuhashi M., 1966; Vollmer & Höltje, 2004). PG constitutes an essential structural component required for cell integrity, and its disruption leads to bactericidal cell lysis (Vollmer *et al*, 2008), making it a major target for antimicrobials (Alderwick *et al*, 2015). However, the unique structure of mycobacterial PG and the alarming emergence of resistance strategies render many of these antimicrobials ineffective against mycobacteria (Catalão *et al*, 2019; Batt *et al*, 2020). For *Mtb*, this pathogen is considered naturally resistant to most β-lactam antibiotics due to the highly active β-lactamase (BlaC), which neutralizes many β-lactams, and the presence of non-classical L,D-transpeptidases, which are inherently resistant to these antibiotics antibiotics (Batt *et al*, 2020; Catalão *et al*, 2019). TB and other mycobacterial IDs still represent a major health burden worldwide, which is exacerbated by the increasing emergence of multi-resistant strains, as well as the high toxicity and long-term treatments associated with the drugs currently in use (Catalão *et al*, 2019). This stresses the urgent need for new bacteriostatics, especially in a time marked by an alarming shortage of new antibacterial classes. In this context, bacteriophage endolysins emerge as a novel and promising class of specific antimicrobials (enzybiotics) for treating bacterial IDs, including those caused by mycobacteria (Dams & Briers, 2019). Bacteriophage-encoded endolysins are PG-hydrolases that target the bacterial cell envelope, enabling host lysis and release of viral progeny at the end of the lytic cycle (Dams & Briers, 2019). Phage endolysins are highly specific, and their exogenous application has been shown to lead to a rapid and specific elimination of pathogenic bacteria *in vitro* and in animal models of infection caused by various pathogens, such as *Streptococcus* sp. (Nelson *et al*, 2001; Loeffler *et al*, 2003), *Bacillus anthracis* (Schuch *et al*, 2002) and *Staphylococcus aureus* (Haddad Kashani *et al*, 2018). Moreover, given the essentiality of the PG molecule, endolysins have poor prospects for developing resistance, further supporting enzybiotics as an attractive alternative to antibiotics (Dams & Briers, 2019).

Phages infecting mycobacteria (mycobacteriophages) produce an endolysin protein referred to as Lysin A (LysA), whose lytic action is assisted by a second enzyme with esterase activity, referred to as Lysin B (LysB), which cleaves the bonds connecting MAs and AG molecules molecules (Catalão *et al*, 2019; Dams & Briers, 2019). The bactericidal activity of mycobacteriophage endolysins has been reported for LysA from the *PDRPxv* mycobacteriophage, capable of lysing the non-pathogenic *Mycobacterium smegmatis* (*Msm*), a lytic effect that was enhanced by its combination with *PDRPxv*LysB (Eniyan *et al*, 2020). More recently, an enzyme cocktail combining LysA and LysB lysins with capsule-degrading enzymes has exhibited bactericidal effects on several pathogenic species of mycobacteria (Bartlett *et al*, 2024), which reinforces the potential of mycobacteriophage LysA endolysins as antimicrobials, alone or in combination with other lysins and drugs (Eniyan *et al*, 2020; Bartlett *et al*, 2024). Thus, a detailed understanding of the mechanisms underlying PG binding and hydrolysis by LysA endolysins at a molecular level is key to their rational development and application as effective enzybiotics. To this end, bioinformatic studies of LysA endolysins have revealed them to be complex multi-domain enzymes. Predominantly, they exhibit a three-domain arrangement formed by N-terminal and central catalytic domains with putative peptidase, glycoside-hydrolase, and/or amidase activities, followed by a C-terminal region reported to function as a PG-binding domain in *D29*LysA (Gangakhedkar & Jain, 2024; Payne & Hatfull, 2012). This complex architecture aligns with the intricate structure of mycobacterial PG, a complexity that has likely hampered their structural and functional analysis so far, as evidenced by the scarce experimental structural information available, with, to our knowledge, only computational models reported for LysA from the bacteriophages *TM4* (Urdániz *et al*, 2022), *Che12* (Saadhali *et al*, 2016), *D29*LysA (Joshi *et al*, 2017, 2019; Gangakhedkar & Jain, 2024).

Among known mycobacteriophages, *D29* and *DS6A* are able to infect pathogenic species of mycobacteria, including *Mtb* (W.B. Redmond & J. C. Carter, 1960; Yang *et al*, 2024; Froman *et al*, 1954). Based on this information, LysA endolysins encoded by *D29* and *DS6A* phages are expected to efficiently cleave the PG of *Mtb* and other pathogenic mycobacteria, thus emerging as relevant candidates for enzybiotic development targeting their infections. Both *DS6A*LysA (residues 1-486) and *D29*LysA (residues 1-493) exhibit a three-domain architecture (**Fig. 1**) that, in the case of *D29*LysA, consists of an N-terminal N4 domain with proposed cysteine protease activity, followed by a glycosyl hydrolase domain belonging to family 19 (referred to as the GH19 domain), and a C1 domain reported to interact with the PG molecule (Payne & Hatfull, 2012; Gangakhedkar & Jain, 2024). *DS6A*LysA is comprised of an N-terminal GH19 domain followed by an amidase domain predicted to be a member of the Amidase_2 family (referred to as Ami2B domain), and a C2 suggested to function as a PG-binding region (Payne & Hatfull, 2012). Biochemical studies conducted on *D29*LysA endolysin show that *D29*N4 and *D29*GH19 are indeed domains exhibiting PG-hydrolase activity, where *D29*N4 is required for optimal bacterial lysis (Joshi *et al*, 2017; 2022; Gangakhedkar & Jain, 2024). Moreover, analysis of *D29*N4, combining computational modeling with thorough biochemical characterization, has recently evidenced the proposed peptidase function predicted for this domain (Gangakhedkar & Jain, 2024). However, the lack of experimental structures to date has prevented the validation of *in silico* models available for *D29*N4, as well as other homologous GH19 and Ami2B domains (Joshi *et al*, 2017; Saadhali *et al*, 2016; Urdániz *et al*, 2022; Nair & Jain, 2024; Gangakhedkar & Jain, 2024), preventing the acquisition of detailed insights into their mechanism of action at a molecular level. This information is highly relevant for future investigations exploring the potential of *DS6A*LysA and *D29*LysA as enzybiotics.

In this work, we present an extensive structural and functional study aimed at the characterization of the catalytic domains of *D29*LysA and *DS6A*LysA endolysins using X-ray crystallography, small-angle X-ray scattering, in combination with site-directed mutagenesis and molecular docking approaches. To our knowledge, our results constitute the first experimental structures reported for mycobacteriophage endolysins, which unveil key aspects of their mode of interaction with PG and hydrolysis. This work provides a solid foundation for the future therapeutic application of *D29*LysA and *DS6A*LysA, including the rational design of optimized enzybiotics to fight mycobacterial infections.

## 2. Material and methods

### 2.1. Protein production

Protein construct bearing the catalytic domains of *D29*LysA (UniProt entry code O64203) and *DS6A*LysA (UniProt entry code G8I4E0) endolysins were designed for their recombinant production. These include each individual wild-type domain *D29*N4 (residues 1-174), *D29*GH19 (residues 179-370), *DS6A*GH19 (residues 1-196), and *DS6A*Ami2B (residues 206-395), the catalytically inactive mutant *D29*GH19^E228Q^ (designed based on available activity studies of GH19 family (Hoell *et al*, 2006), and a construct bearing both GH19 and Ami2B domains of *DS6A*LysA, (*DS6A*GH19-Ami2B, residues 1-395). Constructs were designed as N-terminal His-tagged proteins followed by a 3C-protease site preceding *D29*LysA and *DS6A*LysA catalytic domains. The DNA coding for the constructs cloned into a pET-28a plasmid (between NcoI and XhoI restriction sites) was purchased to GenScript with codon optimization for *E. coli* expression. Proteins were produced as recombinant proteins in *E. coli* BL21 (DE3) Star (Invitrogen) using 2xTY media supplemented with kanamycin (50 μg/ml). Cells were grown at 37 ºC until an OD_600 nm_ of 0.8 and then cooled down to 16 ºC for an hour. Expression was then induced at 16 ºC by the addition of 1 mM IPTG for 20 h. Cells were harvested by centrifugation at 5251 *g* for 20 min at 4 ºC, and cell pellets were stored at −20 ºC.

Cell pellets were thawed and resuspended in 20 mM Tris-HCl, 500 mM NaCl, 10 mM imidazole (buffer A), supplemented with 1 mM MgCl_2_, 1 mM MnCl_2_ and 1 µg/ml DNAse (Sigma). This step was performed at RT while subsequent purification steps were carried out at 4 ºC. Cells were disrupted by sonication and the clarified extract was loaded into a 1 ml HisTrap HP crude column (Cytiva) previously equilibrated in buffer A. Protein samples were eluted using an imidazole gradient (from 10 mM to 500 mM) in buffer A, and then buffer exchanged to 20 mM Tris-HCl, 150 mM NaCl (Buffer B) using a PD-10 desalting column (Cytiva). When required, samples were diluted to 1 mg/ml in buffer B prior to overnight cleavage with the 3C protease. Cleaved and not cleaved proteins were concentrated using Amicon 10 and 30 KDa MWCO concentrators (Millipore), and further purified by size exclusion chromatography (SEC) using a Superdex 200 (16/60) or a Superdex 200 10/300 Increase GL column (Cytiva) preequilibrated in buffer B. Samples corresponding to *DS6A*GH19, *DS6A*Ami2B, *DS6A*GH19-Ami2B, *D29*N4 and *D29*GH19 (WT and mutant) were purified using buffers A and B at pH 8.0 supplemented with 4 mM βME (IMAC steps) and 1 mM DTT (SEC). Pure protein fractions obtained from SEC were pooled, concentrated, and flash-frozen in liquid nitrogen for their storage at −80 ºC. Protein sample purity and quantification were assessed using PAGE-SDS and UV-Vis absorbance at 280 nm, respectively.

### 2.2. Crystallisation and structure determination

All crystallization experiments were carried out using the sitting-drop vapour diffusion method at 18ºC in 96-well MRC plates (Hampton Research) and employing an Oryx8 robot (Douglas Instruments). Crystals of *D29*N4, *D29*GH19^WT^, *D29*GH19^E228Q^, *DS6A*GH19, and *DS6A*Ami2B were initially grown in native conditions using the untagged proteins in all cases but for the *D29*N4 domain. *D29*N4 crystallised in 300 nl droplets formed by mixing 150 nl of protein at 11 mg/ml with 150 nl of precipitant solution (0.1 M HEPES pH 7.5, 1.8 M NaCl). *D29*GH19^WT^ and *D29*GH19^E228Q^ crystallised in up to four crystal forms (I-IV). Crystal form I grew in 200 nl droplets formed by mixing 100 nl of *D29*GH19^WT^ at 8.25 mg/ml with 100 nl of precipitant solution (12% (v/v) ethylene glycol; 6% (w/v) PEG 8K, 0.1 M imidazole/MES buffer pH 6.5, and DL-amino-acids (glutamic acid, alanine, glycine; lysine, serine) at 20 mM); crystal form II grew in 200 nl droplets formed by mixing 100 nl of *D29*GH19^WT^ at 8.25 mg/ml with 100 nl of precipitant solution (9% (v/v) MPD; 9% (v/v) PEG 1K; 9% (w/v) PEG 3350. 0.1 M imidazole/MES buffer pH 6.5, and DL-amino-acids (glutamic acid, alanine, glycine; lysine, serine) at 20 mM); crystal form III grew in 200 nl droplets formed by mixing 100 nl of *D29*GH19^WT^ at 8.25 mg/ml with 100 nl of precipitant solution (9% (v/v) MPD, 9% (w/v) PEG 1K, 9% (w/v) PEG 3350, 0.1 M imidazole/MES buffer pH 6.1, and DL-amino-acids (glutamic acid, alanine, glycine; lysine, serine) at 20 mM; or 13% (v/v) MPD, 9% (v/v) PEG 1K, 9% (w/v) PEG 3350, 0.1 M imidazole/MES buffer pH 6.1, and DL-amino-acids (glutamic acid, alanine, glycine; lysine, serine) at 20 mM; crystal form IV grew in 200 nl droplets formed by mixing 100 nl of *D29*GH19^WT^ at 8.25 mg/ml with 100 nl of precipitant solution (9% (v/v) MPD, 9% (w/v) PEG 1K, 9% (w/v) PEG 3350, 0.1 M Tris/BICINE pH 8.5, 30 mM MgCl_2_, 30 mM CaCl_2_); and crystal form V grew in 300 nl droplets formed by mixing 100 nl of *D29*GH19^E228Q^ at 9 mg/ml with 200 nl of precipitant solution (20 % (v/v) PEG 500 MME, 10 % (w/v) PEG 20K, 0.1 M HEPES/MOPS buffer pH 7.5, and DL-amino-acids (glutamic acid, alanine, glycine; lysine, serine) at 20 mM. *DS6A*GH19^WT^ crystallised in 300 nl droplets formed by mixing 150 nl of protein at 11 mg/ml with 100 nl of precipitant solution (2.2 M NaCl, Tris-HCl 50 mM pH 8.0), and 50 nl of a seed stock in stabilised in the precipitant solution with the butter at 100 mM. *DS6A*Ami2B^WT^ crystallised in 300 nl droplets formed by mixing 150 nl of protein at 16.5 mg/ml with 150 nl of precipitant solution (0.8 M NaH_2_PO_4_, 0.4 M KH_2_PO_4_ and 0.1 M HEPES at pH 7.5). Complexes of *D29*GH19^WT^:NAG and *D29*GH19^WT^:NAG_2_:NAG were obtained by soaking native *D29*GH19^WT^ crystals into a precipitant solution supplemented with 31 mM NAG_6_ for 30 min and 10 seconds, respectively. The *D29*GH19^E228Q^:NAG_4_ complex was obtained by *in situ* soaking, in which NAG_4_ in powder was added to the crystallisation drop containing native *D29*GH19^E228Q^ crystals that were incubated for 45 min. A complex of *DS6A*Ami2B^WT^ bound to the PG fragment L-Ala-D-*iso*-Gln-L-Lys-D-Ala-D-Ala, here referred to as P5^Lys^ (*DS6A*Ami2B^WT^:P5^Lys^ complex) was obtained by soaking native crystals of *DS6A*Ami2B in the precipitant solution supplemented with 100 mM of P5^Lys^ for 30 s. NAG_4_, and NAG_6_ were purchased to Megazyme and the P5^Lys^ peptide was purchased to GenScript. Crystals were flash-cooled in liquid nitrogen without cryoprotection for X-ray data collection, which enabled to measure complete X-ray data sets at 100*K* using synchrotron radiation at ALBA (Barcelona, Spain) and at ESRF (France). Data sets were integrated, scaled and reduced using XDS (Kabsch, 2010) and AIMLESS (Evans & Murshudov, 2013). Data collection statistics are summarized in Table 1SM. Structure resolution was carried out using the molecular replacement method with PHASER (McCoy *et al*, 2007). For native *D29*N4^WT^, *D29*GH19^WT^, *DS6A*GH19^WT^ and *DS6A*Ami2B structures, the coordinates corresponding to models of each individual domain predicted with AlphaFold2 (Evans *et al*, 2022; Jumper *et al*, 2021)were used as search models. For remaining structures, the experimental coordinates of *D29*GH19^WT^ and *DS6A*Ami2B were used as search models. A single and unambiguous solution was obtained in all cases. Initial models were subjected to alternate cycles of model building with COOT (Emsley *et al*, 2010), and refinement with PHENIX (Adams *et al*, 2010) and BUSTER (Bricogne G. *et al*, 2017). NCS restrictions were used during the refinement of structures with more than one protein monomer in the asymmetric unit. Regularised coordinates of the P5^Lys^ molecule were generated with MERCURY (MacRae *et al*, 2020). The geometry of the final models was validated using MOLPROBITY (Chen *et al*, 2009). Figures were generated using PYMOL (Version 3.1; Schrödinger). The atomic coordinates and structure factors have been deposited in the Protein Data Bank under the accession codes indicated in Table 1SM.

### 2.3. Sequence and tertiary structure alignment

Comparative analysis of *D29*N4, *D29*GH19, *DS6A*GH19, and *DS6A*Ami2B domains with homologue domains was performed using sequence alignment with Clustal Omega Multi Sequence Alignment (Sievers *et al*, 2011). Superimposition of three-dimensional structures was carried out using PYMOL (Version 3.1; Schrödinger). Search for structurally homologue domains of *D29*N4 and *DS6A*Ami2B were carried out using the DALI server (Holm *et al*, 2023).

### 2.4. Molecular docking with PG fragments

A complex of *D29*GH19^WT^ with a peptidoglycan fragment was modelled by molecular docking using the GOLD (Genetic Optimization for Ligand Docking) software (version 2024.3.0) (Verdonk *et al*, 2003). The PG fragment consisted on four saccharide units linked to a dipeptide moiety (NAG-NAM°[L-Ala-D-*iso*-Gln])_2_ (4S2P), thus resulting in the complex referred to as *D29*GH19:4S2P. The coordinates of the 4S2P ligand were extracted from a Gram-negative-PG model provided by Prof. Shahriar Mobashery (Meroueh *et al*, 2006), and formatted with MERCURY (MacRae *et al*, 2020). GOLD uses the Genetic algorithm (GA) method that allows a partial flexibility of the protein and full flexibility of ligand. Thirty independent GA runs, a maximum of 100.000 GA operations, were performed on a set of five groups with a population size of 100 individuals. An ensemble docking was performed using the four independent chains found in the *D29*GH19^WT^:NAG_2_:NAG crystal structure and a PG-fragment ligand, with a search efficiency of 200 % (very flexible docking). All water molecules and ligands were removed from the protein. Finally, the CHEMPLP scoring function was used with default parameters to evaluate the solutions.

A model of *DS6A*LysA catalytic domains (*DS6A*GH19-Ami2B) interacting with the bacterial PG was manually generated by superimposing the experimental complexes *D29*GH19^E228Q^:NAG_4_ and *DS6A*Ami2B:P5^Lys^ onto the SAXS model using PyMol (Version 3.1; Schrödinger). The resulting model was next correlated with a larger Gram-negative PG scaffold reported by NMR (Meroueh *et al*, 2006).

### 2.5. Small-angle X-ray scattering (SAXS)

SAXS experiments were performed at the beamline B21 of the Diamond Light Source (Harwell Campus, UK) (Cowieson *et al*, 2020). Samples of 45 µl of 6His-tagged and untagged *DS6A*GH19-Ami2B protein at 5.9 mg/ml were loaded into a SRT-C SEC-300 (Sepax) column equilibrated in buffer (20 mM Tris pH 8.0, 150 mM NaCl and 1 mM DTT) and connected to an Agilent 1200 HPLC system at 18 °C. Continuously eluting samples were exposed for 3 s in 10 s acquisition blocks using an X-ray wavelength of 1 Å, and a sample to detector (Eiger 4M) distance of 3.7 m. The data covered a momentum transfer range of 0.0032 < q < 0.34 Å^−1^. The frames recorded immediately before elution of the sample were subtracted from the protein scattering profiles. The Scåtter software package (www.bioisis.net) was used to analyse data, buffer-subtraction, scaling, merging and checking possible radiation damage of the samples. The R_g_ value was calculated with the Guinier approximation assuming that at very small angles q < 1.3/R_g_. The particle distance distribution, D_max_, was calculated from the scattering pattern with GNOM, and shape estimation was carried out with DAMMIF/DAMMIN, all these programs included in the ATSAS package (Manalastas-Cantos *et al*, 2021). The proteins molecular mass was estimated with GNOM. Interactively generated PDB-based homology models were made using the program COOT (Emsley *et al*, 2010) by manually adjusting the X-ray structures obtained in this work, into the envelope given by SAXS until a good correlation between the real-space scattering profile calculated for the homology model matched the experimental scattering data. This was computed with the program FoXS (Schneidman-Duhovny *et al*, 2016).

## 3. Results and discussion

### 3.1. *D29*N4 exhibits a cysteine peptidase-domain belonging to the C40 protease family

#### 3.1.1. Crystal structure of the *D29*N4 domain

The *D29*N4 structure consists of a papain-fold like core formed by a central six-stranded antiparallel β-sheet (β1’-β1-β5) flanked by one α-helix (α2) (**Fig. 2A**). This core is flanked, on one side, by three additional helices (α1, α3 and α3’) neighbouring α2 and the loops interconnecting the four helices. A long-loop (β4-β5 loop) enfolds the β-sheet on the other side. The β-sheet accommodates two additional loops connecting β1-β2 and β3-β4 strands. This structure is conserved in both molecules integrating the asymmetric unit (RMSD of 0.218 Å). A structural homology search across the Protein Data Bank using DALI server revealed *D29*N4 to exhibit homology with members of the NlpC/P60 large family of cell-wall related cysteine peptidases belonging to the peptidase family C40 (MEROPS data base). The latter include dipeptidyl-peptidases hydrolysing peptidoglycan peptide stem between L-Ala-×-D-*iso*-Glu and L-Zaa-Yaa (where Zaa corresponds to L-lysine or *m*DAP; and Yaa is D-Ala or D-Ala-D-Ala) such as the autolysin Acd24020 from *Clostridium difficile* (*Cdf*Acd24020, PDB entry 7CFL) (Sekiya *et al*, 2021), YkfC endopeptidase from *Bacillus cereus* (*Bcr*YkfC, PDB entry 3H41) (Xu *et al*, 2010), and two RipA and RipB peptidoglycan hydrolases from *Mtb* (*Mtb*RipA and *Mtb*RipB; PDB entry codes 3PBC and 3PBI, respectively) (Böth *et al*, 2011). This aligns with the PG-hydrolase activity initially predicted for this domain (Payne & Hatfull, 2012), and with recent studies by Gangakhedkar and Jain, where the authors map key residues for *D29*N4 activity using site-directed mutagenesis based on sequence homology studies and computational modelling of *D29*N4 (Gangakhedkar & Jain, 2024).

**Figure 2.**
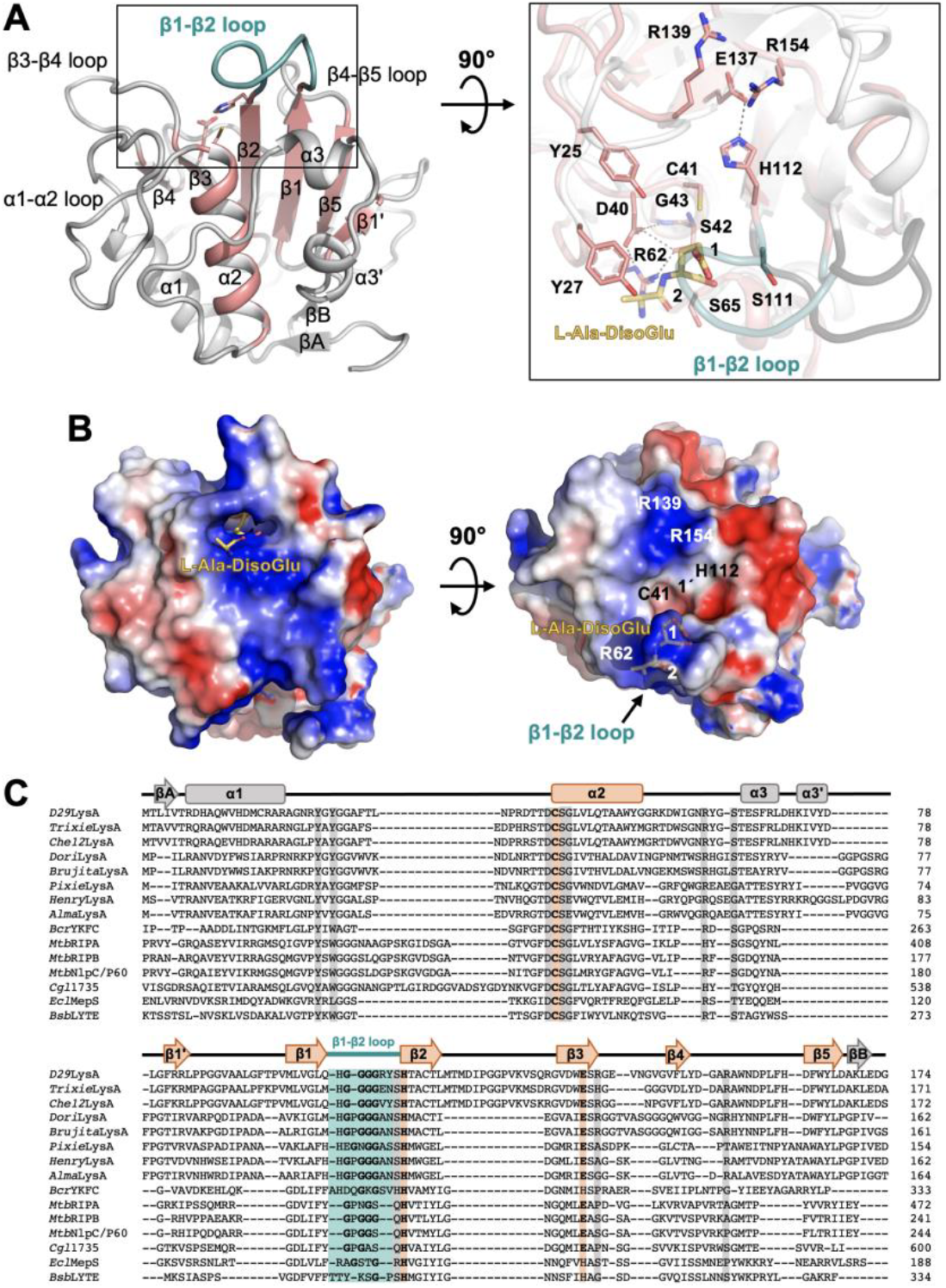
(A) General representation of the *D29*N4 crystal structure as cartoons with the papain-like core fold highlighted in salmon. Loops forming the PG-binding site are labelled and the β1-β2 loop is emphasised in colour in blue (left). Top view of the PG-binding channel of *D29*N4 (in salmon) superimposed with the structure of the *Bcr*YkfC:L-Ala-D-iso-Glu complex (in grey) (right). *D29*N4 residues proposed to participate in PG biding and hydrolysis are represented as sticks and superimposed. The PG fragment corresponding to the *Bcr*YkfC:L-Ala-D-iso-Glu complex is depicted as yellow sticks. (B) Representation of the electrostatic potential surface of *D29*N4 showing PG fragment bound at the *Bcr*YkfC:L-Ala-D-iso-Glu complex that is superimposed and depicted as yellow sticks. Details of the catalytic dyad (C41 and H112) and arigines proposed to participate in PG interaction across positions 2, 1 and 1’ are labelled. (C) Sequence alignment of *D29*N4 with homologue peptidase domains found at LysA endolysins from the mycobacteriophages *Trixie* (*Trixie*LysA; UniProt entry G1JV62), *Che12* (*Che12*LysA; UniProt entry Q1A0K6), *Brujita* (*Brujita*LysA; UniProt entry B5U397), *Pixie* (*Pixie*LysA, UniProt entry G1D539), *Henry* (*Henry*LysA, UniProt entry G1D285), *Alma* (*Alma*LysA; UniProt entry G8I7N0); and other homologue domains including *Bcr*YkfC (UniProt entry), *Mtb*RipA (UniProt entry O53168); *Mtb*RipB (UniProt entry P9WHU5), *Mtb*Nlp/P60 (UniProt entry Q740Y8), *Cg*1735 (UniProt entry Q8NQA0), *Rcl*MepS (UniProt entry P0AFV4) and *Bsb*LYTE (UniProt entry P54421).

*The Bcr*YkfC peptidase has been shown to hydrolyse the L-Ala-γ-D-*iso*-Glu-L-Lys tetrapeptide between the glutamate and lysine residues, and its crystal structure has been determined in complex with the reaction product L-Ala-γ-D-Glu (*Bcr*YkfC:L-Ala-γ-D-Glu, PDB entry 3H41) that binds at positions 2 and 1 of the PG-binding channel (Xu *et al*, 2010). The structural comparison of *D29*N4 with *Bcr*YkfC:L-Ala-γ-D-Glu reveals a large and positively charged channel expected to accommodate the PG substrate in *D29*LysA (**Fig. 2AB**). This channel is formed by residues belonging to the α3 helix and the loops α1-α2, α2-α3, β1-β2, β3-β4 and β4-β5, and it accommodates the catalytic triad highly conserved across the C40 family of cysteine peptidases (Griffin *et al*, 2023; Ozhelvaci & Steczkiewicz, 2023). The latter consists, on the one hand of a nucleophilic cysteine residue (C41) located at the N-terminus of α2 and a histidine residue (H112) that acts as the base (**Fig. 2A**). The catalytic role of both residues has been validated by site-directed mutagenesis in *D29*N4 (Gangakhedkar & Jain, 2024) and in its homologue peptidases *Cdf*Acd24020 and *Mtb*RipA (Squeglia *et al*, 2014). Our crystal structure of *D29*N4 shows that the position of the catalytic C41 at the active site is fastened by a network of interactions constituted by D40 and R62, forming a salt-bridge interaction, and their H-bond contacts with S42 and G43 residues, which are conserved in other homologue peptidases (**Fig. 2A**). The third constituent of the catalytic triad in C40 peptidases typically consists of a glutamate, histidine, glutamine or asparagine located at the C-terminus of the β3 strand and adjacent the catalytic histidine (Anantharaman & Aravind, 2003). This residue is conserved in *D29*N4 (E137) as well as in its homologue PG-hydrolases *Mtb*RipA (E444) and *Mtb*RipB (E213). The proposed role of this glutamate is to enhance the nucleophilic character of the catalytic cysteine by properly orienting the catalytic histidine towards its imidazole group (Ozhelvaci & Steczkiewicz, 2023), and it has been shown to be necessary for *D29*N4 activity (Gangakhedkar & Jain, 2024). In *D29*N4, the imidazole moiety of H112 orients favourably towards the C41 and E137 side-chains at H-bond distance, thus supporting the correct configuration of the catalytic triad, and the role proposed for E137. While these features align with the *in silico D29*N4 structure predicted by Gangakhedkar and Jain, their computational model shows deviations with respect to our crystallographic structure at the level of the N-terminus of α2 accommodating the catalytic cysteine. These deviations involve D40 and C41 main-chain carbonyl groups, which are stablishing H-bond contacts with the main-chain nitrogen of L44. This forces a distortion of the helical arrangement of α2 unseen in with our crystal structure, where the α-helix exhibits a canonical network of H-bonds.

Structural comparison of *D29*N4 with other N4 domains and homologue peptidases reveals variances in the loops forming the PG-binding channel. This includes longer connectors regions spanning α2-α3, α3-β1’ and β1-β2 and β2-β3 in N4 domains. Also, a significantly longer β2-β3 loop is observed in only some LysA-N4 domains, including *D29*LysA. These prolonged loops contain glycine residues expected to confer them with flexibility, which is likely to assist on substrate interaction. This is particularly remarkable in the case of the β1-β2 loop, highly rich in glycine residues, which aligns with the high B-factor values observed in this region (**Fig. 2SM**). Interestingly, this loop fold towards the Y27 in *D29*N4, thus closing the substrate-binding channel. Of note, this closed conformation is preserved in both molecules of the crystal asymmetric unit. The superimposition of *D29*N4 and *Bcr*YkfC:L-Ala-γ-D-*iso*-Glu illustrates how the β1-β2 loop of *D29*N4 is adopting a close conformation that must undergo a conformational change in order to unlock the substrate-binding site and enable substrate interaction akin to the one observed in the *Bcr*YkfC:L-Ala-γ-D-*iso*-Glu complex (**Fig. 2A**). This points at the β1-β2 loop as a mobile element with a potential regulatory role in the interaction of *D29*N4 with the peptidoglycan.

Based on the superimposition of *D29*N4 and *Bcr*YkfC:L-Ala-γ-D-*iso*-Glu, we proceeded to inspect those interactions involved in the stabilisation of the L-Ala-γ-D-*iso*-Glu moiety, common in both mycobacterial and Gram-positive peptidoglycan molecules. This shows a conserved patter of contacts comprising hydrophobic (Y25 and Y27), charged (D40, R62), and neutral-polar (S42, G43, S65) residues decorating the subsites 1 and 2 of the PG-binding cavity (**Fig. 2AC**). Importantly, these residues are conserved across homologue peptidases including other LysA-N4 domains; as well as in *Cdf*Acd24020, where they are crucial in substrate interaction (Gangakhedkar & Jain, 2024), thus supporting a similar key role in *D29*N4 (**Fig. 2C** and **Table 2SM**). This aligns with homology and mutagenesis studies reported by Gangakhedkar and Jain, where mutations of Y25A and S111A in *D29*N4 are shown to impair the protein activity (Gangakhedkar & Jain, 2024). Moreover, the superimposition of *D29*N4 and BcrYkfC:L-Ala-γ-D-*iso*-Glu also highlights the presence of two arginine residues (R139 and R154) in *D29*N4 that are likely contributing to the 1’ site accommodating the carboxylate group of the mDAP (**Fig. 2**). This analysis differs from computational studies by Gangakhedkar and Jain, where the authors propose a *D29*N4:PG model predicting the mDAP moiety to orient towards R109 and Y110 residues of the β1-β2 loop, which seems to adopt an open conformation in the model (Xu *et al*, 2010; Gangakhedkar & Jain, 2024). Our sequence analysis shows that while R109 and Y110 are not conserved in the sequence of other LysA-N4 domains (**Fig. 2C**). However, R139 and R154 do exhibit conservation across mycobacteriophage endolysins thus supporting these two arginines as expected interactors of the mDAP moiety, contributing to stabilising the peptide moiety of the PG in a similar fashion to the one proposed for the *Cdf*Acd24020 autolysin (Sekiya *et al*, 2021).

### 3.2. *D29*GH19 and *DS6A*GH19 are flexible “loopless” GH19 domains

#### 3.2.1. General structure of *D29*GH19 and *DS6A*GH19 domains

We have determined the crystallographic structures of the wild-type *D29*GH19 domain (*D29*GH19^WT^) solved in three different crystal forms (*D29*GH19^WT-I^, *D29*GH19^WT-II^ and *D29*GH19^WT-III^) at 1.6-2.0 Å resolution, and the crystal structure of the wild-type *DS6A*GH19 domain (*DS6A*GH19^WT^) at 2.8 Å resolution. *D29*GH19^WT^ and *DS6A*GH19^WT^ exhibit a structure rich in alpha helices that arrange in two lobes (**Fig. 3A**). A first lobe is formed by residues provided by N-terminal (residues 179-233 in *D29*GH19^WT^) and C-terminal (residues 307-370 in *D29*GH19^WT^) regions, referred to as NC-lobe, that comprises helices α1-α4 and α7-α10. The second lobe is built by residues belonging to the middle region of the polypeptide chain (residues 234-308 in *D29*GH19^WT^), here referred to as M-lobe, that comprises helices α6-α7. This fold is conserved in the four structures determined, which is reflected by the RMSD values calculated for: i) *D29*GH19^WT-I^ and *D29*GH19^WT-II^ (0.320 Å for 180 Cα atoms); ii) *D29*GH19^WT-I^ and *D29*GH19^WT-III^ (1.832 Å for 167 Cα atoms) and; *D29*GH19^WT-I^ and *DS6A*GH19^WT^ (1.071 Å for 154 Cα) (**Fig. 3A**). According to (Hoell *et al*, 2006), GH19 domains can be classified as “loopful” or “loopless” according to the presence or absence of insertions in loops I-V and C, located towards the substrate-binding groove, that results in larger or smaller number of substrate-interaction subsites, respectively (Hoell *et al*, 2006). The superimposition of *D29*GH19 with endochitinase C from *Secale cereale* (Rye) (*RSC-c*GH19, PDB entry 4DWX), as a representative of a “loopful” GH19 domain, shows high degree of structural conservation in the two-lobe core structure but clear differences in the length of loops I-V and C, which are much shorter in *D29*GH19 (Fig. 4D and Fig. 4G). This is in line with the sequence alignment of homologue domains found across phages, bacteria and eukaryote, which highlights the presence of short I-V loops in both of *D29*GH19 and *DS6A*GH19 domains, and in GH19 domains present other LysA endolysins (**Fig. 3D**). Recently, GH19 hydrolases have been reclassified in two different subfamilies, referred to as GH19_1 and GH19_2, embracing enzymes with peptidoglycan endo-β-1,4-N-acetylmuramidase and chitin endo/exo-β-1,4-N-acetylglucosaminidase activities, respectively (see <u>CAZY – GH19</u>). Altogether this illustrates LysA-GH19s as “loopless” glycosyl-hydrolases belonging to the GH19_1 subfamily, exhibiting smaller substrate-binding channels when compared to their “loopful” homologues belonging to the GH19_2 subfamily (Hoell *et al*, 2006).

**Figure 3.**
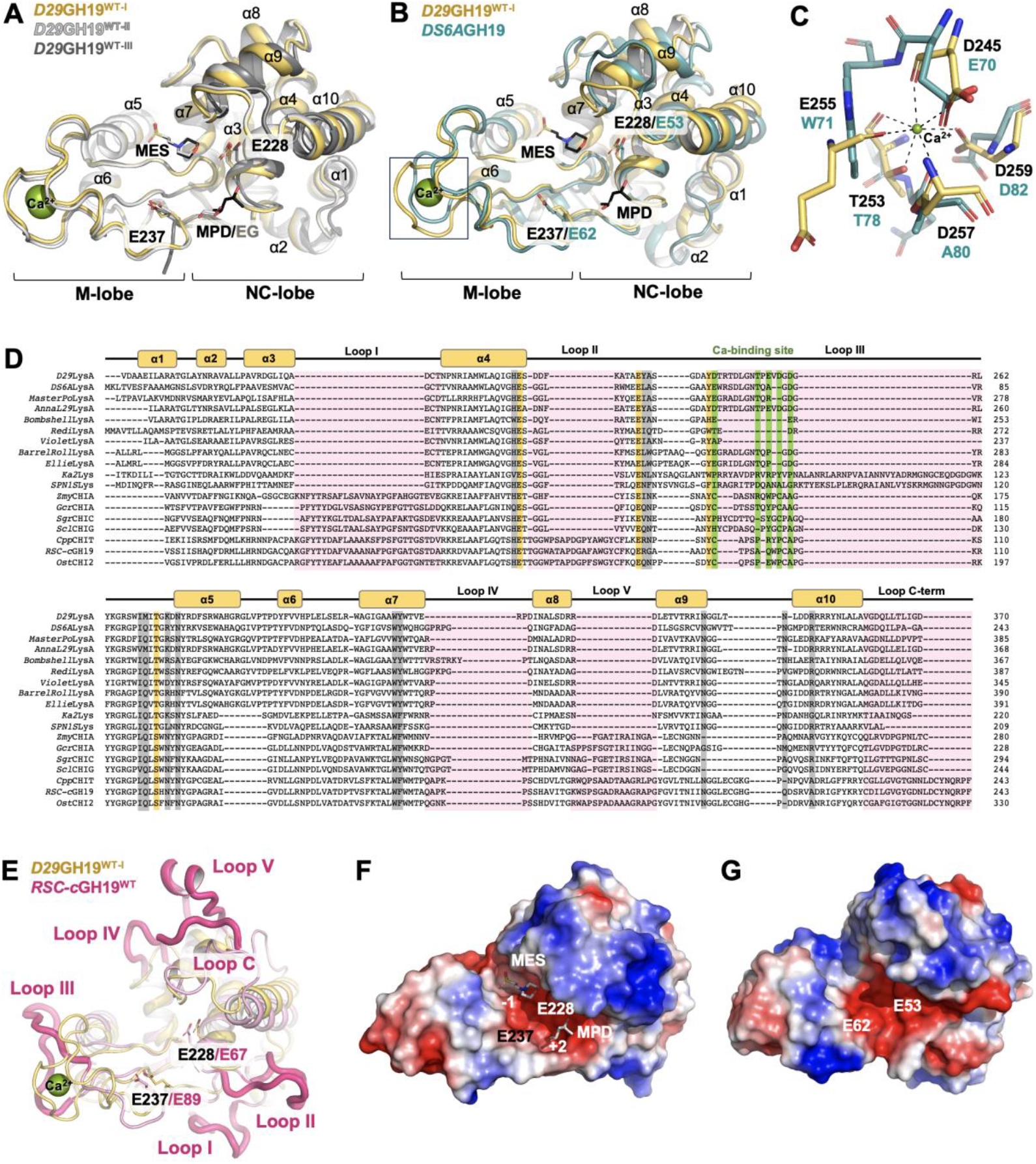
General structure of *D29*GH19 and *D29*GH19 domains. (A) Superimposition of *D29*GH19^WT-I^ (yellow), *D29*GH19^WT-II^ (light grey) and *D29*GH19^WT-III^ (dark grey) crystal structures depicted as cartoons. The catalytic residues and bound molecules coming from the crystallisation condition are represented as sticks, and calcium cations are shown as green spheres. MES and MPD molecules bound to *D29*GH19^WT-I^ are coloured in black; and MES and EG molecules bound to *D29*GH19^WT-II^ are coloured in light grey. (B) Superimposition of *D29*GH19^WT-I^ (yellow) and *DS6A*GH19^WT^ (blue) crystal structures depicted as cartoons, showing the conserved catalytic dyad as sticks. Calcium cations, MES and MPD are represented as in panel (A). (C) Overlay of the calcium-binding site in *D29*GH19^WT-I^ (yellow) and the corresponding region in *DS6A*GH19^WT^ (blue). (D) Details of the sequence alignment of *D29*GH19 and *DS6A*GH19 with other GH19 domains found across bacteria, eukaryota and phages, including mycobacteriophage LysA endolysins: *RSC-c*GH19 (UniProt entry Q9FRV0), endochitinase from *Carica papaya* (*Cpp*CHIT, UniProt entry P85084), chitinase 2 from *Oryza sativa subsp. japonica* (*Ost*CHI2, UniProt entry Q7DNA1), endochitinase A from *Zea mays* (*Zmy*CHIA, UniProt entry P29022), chitinase A from *Gemmabryum coronatum* (*Gcr*CHIA, UniProt entry A9ZSX9), chitinase A from *Streptomuces griseus* (*Sgr*CHIC, UniProt entry O50152), chitinase G from *Streptomyces coelicolor* (*Scl*CHIG, UniProt entry Q8CK55), endolysin from *Pseudomonas phage Ka2* (*Ka2*Lys, UniProt entry A0A9E7MJ36), endolysin from *Salmonella phage SPN1S* (*SPN1S*Lys, UniProt entry H2D0G4), placed Lysin A from *Mycobacteriophage Redi* (*Redi*LysA, UniProt entry G3M5B9), to be red accelerate displace Lysin A from *Mycobacteriophage BarrelRoll* (*BarrelRoll*LysA, UniProt entry G3MCI0), Lysin A from *Mycobacteriophage Ellie* (*Ellie*LysA, UniProt entry A0A7G8LLY0), Lysin A from *Mycobacteriophage MasterPo* (*MasterPo*LysA, UniProt entry A0A7G8LGT1), Lysin A from *Mycobacteriophage AnnaL29* (*AnnaL29*LysA, UniProt entry S5VWI7), Lysin A from *Mycobacteriophage Bombshell* (*Bombshell*LysA, UniProt entry A0A7G8LLL9), and Lysin A from *Mycobacteriophage Violet* (*Violet*LysA, UniProt entry G3MEJ3). (E) Structural superimposition of the *D29*GH19^WT-I^ and the “loop-full” GH19 structure, corresponding to *RSC-c*GH19 (PDB entry 4DWX), represented as yellow and pink cartoons, respectively, with catalytic glutamates depicted as sticks. Loops I, II, III, IV, V and C of *RSC-c*GH19 are labelled and highlighted as thicker dark-pink loops. (F) and (G) show the electrostatic-potential surface representation of *D29*GH19^WT-I^ and *DS6A*GH19^WT^ domains, respectively. Positively and negatively charged residues are represented in colour blue and red, respectively. Catalytic residues and ligands bound at the substrate-binding groove are labelled.

**Figure 4.**
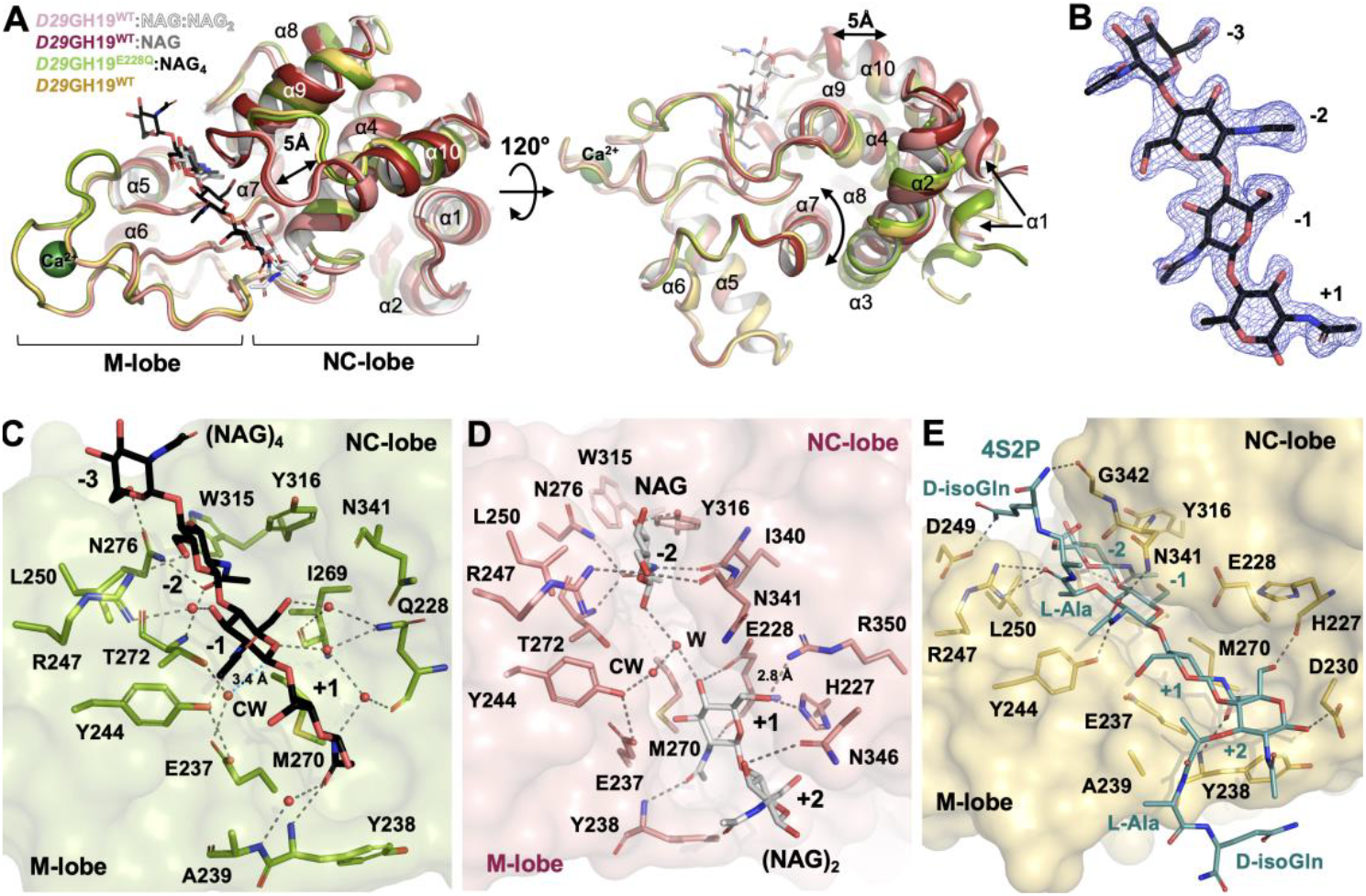
*D29*GH19 complexes with PG analogues. (A) Structural superimposition of *D29*GH19^E228Q^:NAG_4_ and *D29*GH19^WT^:NAG:NAG_2_ complexes with *D29*GH19^WT-I^, represented as cartoons coloured in green, pink and yellow respectively. Bound saccharide molecules are represented as sticks coloured in black (NAG_4_) and light white (NAG and NAG_2_). (B) Detail of the electron density map 2Fo-Fc (blue mesh) corresponding to the NAG_4_ molecule of the *D29*GH19^E228Q^:NAG_4_ complex contoured at sigma = 1.0. (C) and (D) show details of the interaction with NAG_n_ fragments identified in *D29*GH19^E228Q^:NAG_4_ and *D29*GH19^WT^:NAG:NAG_2_ complexes, respectively. H-bonds are highlighted as grey dashed lines. PG analogues and residues involved in ligand binding are represented as sticks following the same colour code as in (A). Water molecules and calcium cations are presented as red and dark green spheres, respectively. (E) Detail of the complex *D29*GH19^WT^:4S2P predicted by docking showing the substrate-binding cavity (protein in yellow). The catalytic residues and the PG fragment are represented as blue and yellow sticks, respectively.

Despite the high structural conservation observed between *D29*GH19 and *DS6A*GH19 models, significant differences are identified in the architecture of the M-lobe as well as subtle deviations in the inter-lobe arrangement. In *D29*GH19^WT^, M-lobe exhibits an EF-hand-like calcium-binding site located at the apical region of the lobe, where the cation appears stabilised by interactions with five amino acids (T253, E255, D245, D257 and D259) resulting in a hepta-coordinated calcium complex (**Fig. 3C**). The presence of a calcium cation at this position is fully supported by a strong signal observed in the 2Fo-Fc electron density map (sigma = 15) that is well interpreted by the refinement of a calcium cation, as well as by the presence at this position of a peak of sigma > 9 in the anomalous electron density map calculated (**Fig. 3SM**). The calcium-interacting site is structurally conserved in both *D29*GH19^WT-I^ and *D29*GH19^WT-II^ structures, but it is not observed in the *D29*GH19^WT-III^ model, where this region is disordered. Interestingly, this calcium-binding site is not present in the *DS6A*GH19^WT^ domain, where three conserved residues (E70, T78 and D82) exhibit a different orientation, and two of the acidic residues interacting with calcium in *D29*GH19^WT^ (E255 and D257) are not conserved (W71 and A80 in *DS6A*GH19^WT^) (**Fig. 3C-D**). To our knowledge, the calcium-binding site present in *D29*GH19 has not been observed in other GH19 structures reported. This is exemplified by the sequence alignment shown in **Fig.3D** that also indicates poor conservation of this calcium-binding site across LysA endolysins (**Fig. 3D**). Superimposition of *D29*GH19^WT-I^ and *D29*GH19^WT-II^ models with *D29*GH19^WT-III^ and *DS6A*GH19^WT^ structures highlights the presence of an inter-lobe shift, which aligns with higher RMSD values calculated (**Fig. 3AB**). In this shift, NC and M-lobes move as independent scaffolds, which agrees with the RMSD values calculated for the individual M-lobes (0.195 Å for 44 Cα atoms) and NC-lobes (0.240 Å for 103 Cα atoms) of *D29*GH19^WT-I^ and *D29*GH19^WT-III^ coordinates; and with the RMSD values calculated for M-lobes (0.57 Å for 57 Cα atoms) and NC-lobes (1.115 Å for 84 Cα atoms) of *D29*GH19^WT-I^ and *DS6A*GH19^WT^ structures. In the case of *D29*GH19, the inter-lobe rotation observed supports an inherent flexibility of this domain relevant for the interaction with the PG substrate, as already proposed in other GH19_1 homologues (Ohnuma et al, 2013).

The substrate-binding site is embedded into a large grove formed by residues belonging to both M and NC-lobes that allocates the catalytic dyad and two glutamate residues (E228/E237 in *D29*GH19; E53/E62 in *DS6A*GH19), in the middle of the cavity (**Fig. 3**). This catalytic dyad is conserved across GH19 members, and their implication in catalysis has been verified through site-directed mutagenesis and biochemical studies in *D29*LysA, as well as the homologue Chitinase G from *Streptomyces coelicolor* and in the bacteriophage *SPN1S* endolysin, reported elsewhere and that we recover in **Table 3SM** (**Fig. 3D**) (Hoell *et al*, 2006; Park *et al*, 2014). Inspection of the electrostatic potential surface of *D29*GH19 and *DS6A*GH19 domains shows along PG-binding channels numerous negative charges, protruding from aspartate/glutamate side-chains as well as main-chain carboxyl groups, which is a typical feature of saccharide-interacting sites/grooves found in carbohydrate-binding domains/hydrolases (**Fig. 3FG**). In *D29*GH19^WT-I^ and *D29*GH19^WT-II^ structures, molecules coming from the crystallisation conditions bind adjacent the catalytic dyad, mimicking saccharide moieties of the substrate at positions −1 (a molecule of MES in *D29*GH19^WT-I^ and *D29*GH19^WT-II^) and +2 (molecules of MPD or ED in *D29*GH19^WT-I^ and *D29*GH19^WT-II^, respectively), thus defining a PG-binding groove with up to three positions (−1, +1 and +2) (**Fig. 3AF**).

#### 3.2.2. *D29*GH19 conserves structural features required for PG hydrolysis and exhibits chitinase activity in *crystallo*

To gain more insights into substrate binding and hydrolysis, crystallographic complexes with NAG_n_ oligosaccharides as glycan analogues were pursued by soaking using the catalytically-active *D29*GH19^WT^ domain and the catalytically inactive mutant E228Q (*D29*GH19^E228Q^) domain. This led us to solve the structure of *D29*GH19^E228Q^ alone and bound to NAG_4_ (*D29*GH19^E228Q^:NAG_4_) at 1.45 Å and 1.83 Å resolution, respectively. Comparison of *D29*GH19^E228Q^ and *D29*GH19^WT^ crystal models indicates high structural conservation (RMSD of 0.118 Å for 176 Cα atoms) (**Fig. 4SM**). The *D29*GH19^E228Q^:NAG_4_ structure consists of one molecule of the complex in the asymmetric unit, in which the saccharide binds from positions −3 to +1 of the PG-binding channel. In addition, the structure of two more complexes were obtained, by soaking *D29*GH19^WT^ crystals into a solution containing NAG_6_, at 2.34 and 3.10 Å resolution (**Fig. 4** and **4SM**). The first one consists of four complexes of *D29*GH19^WT^ bound to NAG and NAG_2_ (*D29*GH19^WT^:NAG:NAG_2_) in the asymmetric unit, where NAG_2_ and NAG are products resulting of NAG_6_ hydrolysis *in crystallo*. The four *D29*GH19^WT^:NAG:NAG_2_ complexes exhibit the same structure (RMSD values between 0.151 Å and 0.264 Å), with saccharide fragments binding with the same pose at positions −2 (NAG) and, +1 and +2 (NAG_2_) of the PG-binding channel. The other complex consists of one *D29*GH19^WT^ domain bound to NAG (*D29*GH19^WT^:NAG) in the asymmetric unit, where the monosaccharide is a product of NAG_6_ hydrolysis *in crystallo* binding at position −2 of the PG-binding site (**Fig. 4** and **4SM**). Superimposition of complexes and native structures shows conservation of the fold across all the models, and the former evidences the capability of *D29*GH19^WT^ to interact with up to five saccharide rings at positions going from −3 to +2 (**Fig. 4A** and **4SM**). Interestingly, superimposition of *D29*GH19^WT^:NAG:NAG_2_ and *D29*GH19^WT^:NAG complexes with native and *D29*GH19^E228Q^:NAG_4_ structures reveals an inter-lobe rearrangement, similar but larger to the one observed between wild-type native structures, resulting in a closure of the PG-binding channel by up to 5 Å in *D29*GH19^WT^:NAG:NAG_2_ and *D29*GH19^WT^:NAG complexes (**Fig. 4AB**). This rearrangement can be explained as a rotation-in-block of the NC-lobe, which pivots towards a central tryptophan (W268) of the M-lobe partially conserved as a tryptophan or a proline in GH19 domains (**Fig. 4** and **4SM**). This movement involves the motion of several hydrophobic residues of helices α3, α4 and α7 that turn over the tryptophan to enable NC-lobe shift. This inter-lobe rearrangement is also observed in the structure of other GH19 chitinases such as *RSC-c*GH19 solved in native and NAG_n_-bound conditions (**Fig. 4SM**), thus showing how this flexibility extends from “loopful” to “loopless” GH19 domains as a general feature of this family of hydrolases.

A more detailed inspection of the PG-binding channel in *D29*GH19^WT^:NAG and *D29*GH19^WT^:NAG:NAG_2_ reveals that the binding mode of the NAG ring at position −2 is conserved in both complexes (**Fig. 4A**). Thus and for simplicity, subsequent structural analysis will focus on *D29*GH19^E228Q^:NAG_4_ and *D29*GH19^WT^:NAG:NAG_2_ complexes. Comparison of the PG-binding channel in these complexes reveals a network of direct or water-mediated interactions participating in saccharide stabilisation that involves several hydrophobic residues (Y238, A239, Y244, L250,I269, M270, W315 and Y316) and polar amino-acids that include the catalytic residues (E/Q228, E237, T272, K274, N276 and N341), provided by both lobes (**Fig. 4C-D**). Interactions stablished via M-lobe are conserved in both *D29*GH19^E228Q^:NAG_4_ and *D29*GH19^WT^:NAG:NAG_2_ complexes, where it is worth highlighting an H-bond network bridging Y244, T272 and the catalytic E237 with a water molecule (W) and the acetyl group of the NAG ring at position −1. These contacts enable the correct orientation of the catalytic base E237 towards the water, thus allowing the CW molecule to locate at a distance (3.4 Å) and orientation to carry out the nucleophilic attach over the C1 atom of the substrate (**Fig. 4**). This H-bond network is observed in structures here reported for the WT and E228Q mutant of *D29*GH19, and other GH19s structures, constituting a structural hallmark of the catalytic mechanism of this family of glycosyl hydrolases (**Fig. 4SM**). The interaction pattern via NC-lobe residues does however vary between *D29*GH19^E228Q^:NAG_4_ and *D29*GH19^WT^:NAG:NAG_2_ structures (**Fig. 4**). This results of the close conformation of the PG-binding channel in *D29*GH19^WT^:NAG:NAG_2_, also observed in the *D29*GH19^WT^:NAG complex. In this close conformation, the side chain of the catalytic E228 shifts towards the hydroxyl group at C4 of the NAG in position +1. This results in a strong H-bonding interaction (distance of 2.8 Å) between both hydroxyl and carboxyl groups compatible with proton exchange during substrate scission (**Fig. 4** and **4SM**). Moreover, the close arrangement of the PG-binding channel results in additional contacts and/or optimisation of pre-stablished interactions of *D29*GH19 with the ligand. On the one hand, there is an approximation Y315 towards the NAG ring in position −2 that strengthens this interaction. On the other hand, the C-terminus of α4, the N-terminus of α10, and the liker connecting helices α9 and α10 shift to pack more tightly against the products, thus resulting in additional contacts via H227, R350, N341 and N346 side chains, and I340 and N341 main-chain carbonyl groups (**Fig. 4D**). These residues are conserved in *DS6A*GH19 but for Y238 and M270 (substituted by a distal tryptophan and a glutamine in *DS6A*GH19, respectively) and a tyrosine (Y98) edging the position −3 of the PG-binding channel in *DS6A*GH19 are missing in *D29*GH19 (D275) (**Fig. 4D** and **4SM**). Our sequence analysis shows high conservation of these amino-acids involved in ligand interaction, but for Y138, A139, K274 in *D29*GH19, that exhibit more divergent variations (**Fig. 4** and **4SM**). Importantly, mutation of several of these residues in *D29*GH19 and lysins *SPN1S* endolysin (reported elsewhere) evidence a clear impact in their interaction with the substrate or its hydrolysis (see **Table 2SM**), which altogether supports their relevance in *D29*GH19 and *DS6A*GH19 performance.

Next, we decided to further explore the mechanism of substrate recognition of a PG fragment by *D29*GH19 using molecular docking approaches. To do so, an unbiased docking search was performed using the close conformation of *D29*GH19^WT^ and the PG-fragment ligand (NAG-NAM[L-Ala-D-*iso*-Gln])_2_ (4S2P). This yielded eleven top solutions (out of thirty) where the PG fragment consistently binds across positions −2 (NAG), −1 (NAM), +1 (NAG) and +2 (NAM), exhibiting a very conserved pose for the glycan strand and discreet variations in the orientation of the peptide moiety (**Fig. 4** and **4SM**). Inspection of the top docking solution, referred to as *D29*GH19^WT^:4S2P complex, shows how the location of the four saccharide rings aligns well with the NAG_n_ molecules interacting with *D29*GH19 *in crystallo*, which supports the reliability of the *in silico* model (see **Fig. 4SM**). The peptide steps protrude from the glycan strand exhibiting a conserved pose at the D-lactyl moieties, which interact with hydrophobic residues at position −1 (Y224 and I250) and position +2 (A239) via their methyl groups, and form H-bond contacts through the carbonyl group at position −1 with R247. L-Ala and D-*iso*-Gln residues of the ligand do show divergent orientations sustained by plausible interactions with residues NC- and/or M-lobes (see **Fig. 4E**). Importantly, the model shows the only glycan arrangement enabling the positioning of the peptide stems without forming clashes with the protein, as a shift of the NAG-NAM saccharides in the channel would generate collisions between the peptide steps pending from C3 carbon of the NAM units and the enzyme. While this further supports the likelihood of our computational supports, it also features the *D29*GH19 domain as an enzyme with muramidase, that not glycosaminidase, activity as the reducing saccharide after hydrolysis at position −1 would be a NAM residue and not NAG. To our knowledge, our studies provide the first evidence supporting the GH19_1 subfamily of hydrolases to function as muramidases cleaving the β-(1,4)- glycosidic bond connecting NAM and NAG saccharides of the PG.

### 3.3. *DS6A*Ami2B consists of a catalytically competent PG-amidase domain

#### 3.3.1. *DS6A*Ami2B exhibits catalytic features characteristic of the Amidase_2 family

We have solved the crystal structure of *DS6A*Ami2B in native conditions, which consists of three polypeptide chains in the asymmetric unit. All of them exhibit the same fold (RMSD of 0.086-0.118 Å for >150 Cα) consisting of a central beta sheet built by two antiparallel (β1 and β4) and four parallel beta strands (β2, β3, β5 and β6) flanked by five alpha helices (α1-α5) (**Fig. 6A**). Structural homology analysis using DALI indicates highest homology with members of the Amidase 2 family, including the cell-wall remodelling amidases from *Staphylococcus aureus* AmiA (*Srs*AmiA) (PDB entry 4KNK; RMSD 3.194 for 36 Cα atoms) (Büttner et al, 2014) and AmpDh2 (*Srs*AmpDh2) (PDB entry 4BJ4; RMSD 0.996 for 60 Cα atoms) (Martínez-Caballero *et al*, 2013), or the amidase AmiD from *E. coli* (*Ecl*AmiD) (PDB entry 2WKX; 1.168A for 62 Cα atoms) (Kerff *et al*, 2010), which aligns with previous homology predictions reported (Urdániz *et al*, 2022). Members of the Amidase 2 family consist of zinc-dependent hydrolases exhibiting *N*-acetylmuramoyl-L-Alanine amidase activity (EC:3.5.1.28), and that cleave the PG molecule between the D-lactyl group of the NAM residue and the first L-Ala amino-acid of the peptide stem (Büttner *et al*, 2015). Activity that has been demonstrated for homologues including the Ami2B domain of the LysA encoded by the mycobacteriophage TM4 (*TM4*LysA) (Urdániz *et al*, 2022), the prophage Ba02 Ba02 PlyL endolysin encoded by *Bacillus anthracis* (*Bnt*PlyL) (Low *et al*, 2005), AmiE autolysin from *Staphylococcus epidermis* (*Spd*AmiE) (Zoll *et al*, 2010) and LytA autolysin from *Streptococcus pneumoniae* (*Spn*LytA) (Mellroth *et al*, 2014; Sandalova *et al*, 2016).

Inspection of the structure of *DS6A*Ami2B reveals a large T-shape channel located at the C-terminus of the beta-sheet core formed by loops β2-α2, α2-β3, β4-α3, β5-α4, and β6-α5 (**Fig. 5B**). The latter constitutes the substrate-binding site that can be described as a transversal glycan-binding groove exhibiting a negatively charged core, and a perpendicular channel involved in the interaction with the peptide moiety and rich in positively charged residues. In line with our homology analysis, a zinc cation locates at the centre of this channel, where it coordinates to two histidines (H251 and H364) and one aspartate (D376) located at the loop β6-α5 and the C-terminus of β2. These residues are highly conserved across Amidase_2 members, and they have been shown to be essential for the activity of members of this family of amidases including *TM4*LysA (Urdániz *et al*, 2022), *Spd*AmiE (Zoll *et al*, 2010), and *Spn*LytA (Mellroth *et al*, 2014; Sandalova *et al*, 2016) (**Fig. 5C** and **Table 4SM**). In *DS6A*Ami2B structure, the metal is also interacting with a phosphate molecule, present in the crystallization condition, resulting in a tetrahedral coordination of the zinc. The proposed role of the zinc is to polarize the water molecule meant to perform the nucleophilic attack onto the scissile bond of the substrate (Büttner et al, 2014; Sandalova et al, 2016). In addition, a catalytic dyad has been proposed to assist the zinc during PG hydrolysis (Zoll *et al*, 2010; Mellroth *et al*, 2014; Sandalova *et al*, 2016). This dyad consists of a glutamate residue located at the C-terminus of β5, which would act as an acid donating a proton to the newly formed N-terminus of the peptide (Zoll *et al*, 2010; Mellroth *et al*, 2014; Sandalova *et al*, 2016). And a histidine charged residue, located two residues upstream the zinc-coordinated aspartate, and proposed to participate in the stabilization of transition state (Zoll *et al*, 2010; Mellroth *et al*, 2014; Sandalova *et al*, 2016). Both residues are conserved in *DS6A*Ami2B (E317 and K374), which lays adjacent the catalytic zinc orienting its side chain towards the cofactor. Altogether these findings support that *DS6A*Ami2B is a catalytically competent amidase exhibiting conserved features of the Amidase 2 family (**Fig. 5**) (Büttner *et al*, 2015). This is well illustrated by the sequence and structural conservation of *DS6A*Ami2B with other homologue amidases depicted in **Fig. 5 and 5SM**. The latter also highlights the fourteen-residues long β5-α4 loop flanking the PG-binding channel. This long loop resembles the one present in other homologues such as *Srs*AmpDh2, where it has been shown to act as a PG-fastening element (Martínez-Caballero *et al*, 2013). Interestingly, and despite differences in sequence, the length of this loop is conserved across other LysA-Ami2B domains that frequently exhibit an aromatic residue located at the middle of the loop, which emerges as a plausible feature participating in PG interaction (**Fig. 5C**).

**Figure 5.**
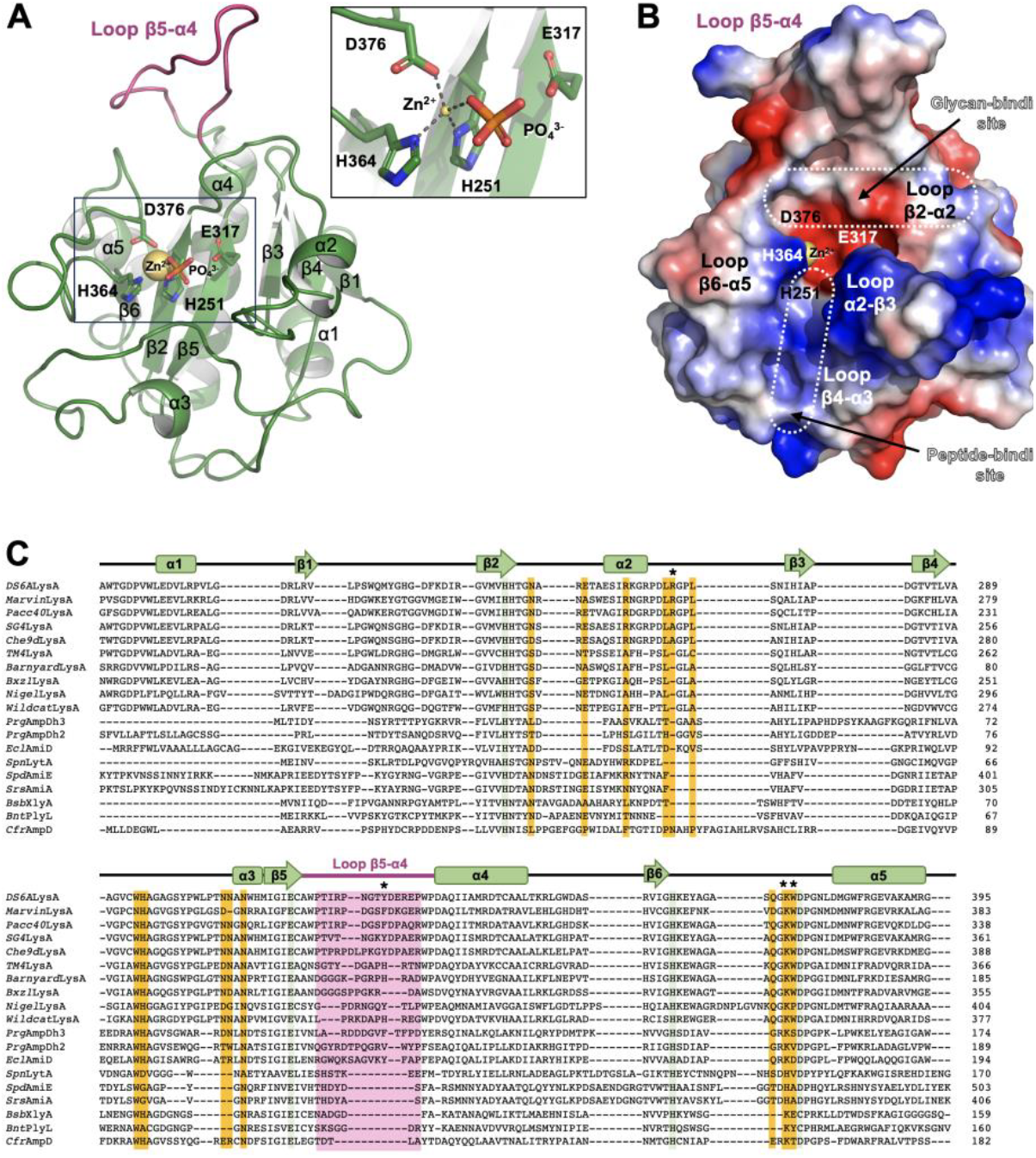
(A) Crystal structure of *DS6A*Ami2B domain represented as green cartoons and highlighting the loop β5-α4 loop in colour purple. Details of the active site are shown on the right box. Catalytic zinc and residues are depicted as a yellow sphere and sticks, respectively. A phosphate molecule found coordinating to the catalytic zinc is presented as sticks. (B) Electrostatic-potential surface representation of *DS6A*Ami2B with the loops contributing to form the PG-binding channel, and the zinc (yellow sphere) and residues involved in catalysis indicated with labels. (C) Details of the sequence alignment of *DS6A*Ami2B with other Amidase_2 domains: AmiA from *Staphylococcus aureus* (*Srs*AmiA; UniProt entry Q2FZK7), AmiE from *Staphylococcus epidermis* (*Spd*AmiE; UniProt entry O33635), LytA from *Streptococcus pneumoniae* (*Spn*LytA; UniProt entry P06653), PlyL from *Bacillus anthracis* (*Bnt*PlyL, UniProt entry A0A6H3AMF3), XlyA from *Bacillus subtilis* (*Bsb*XlyA; UniProt entry P39800), AmiD from *Escherichia coli* (*Ecl*AmiD; UniProt entry P75820), AmpDh2 from *Pseudomonas aeruginosa* (*Prg*AmpDh3; UniProt entry Q9I5D1), AmpDh2 from *Pseudomonas aeruginosa* (*Prg*AmpDh2; UniProt entry Q9HT86), AmpD from *Cytrobacter freundii* (*Cfr*AmpD; UniProt entry P82974), and LysA endolysins from the mycobacteriophages *Stinger* (*Stinger*LysA; UniProt entry G8I9G3), *Nigel* (*Nigel*LysA; UniProt entry B3VLX1), *Bxz1* (*Bxz1*LysA; UniProt entry Q852W9), *Pacc40* (*Pacc40*LysA; UniProt entry B5U576), *SG4* (*SG4*LysA; UniProt entry G8I9P6), *Che9d* (Che9dLysA; UniProt entry Q855S8), *Barnyard* (*Barnyard*LysA; UniProt entry Q856D3), *TM4* (*TM4*LysA; UniProt entry Q9ZX49), *Corndog* (*Corndog*LysA; UniProt entry Q856M5), *Marvin* (*Marvin*LysA; UniProt entry G1BNC1), *Wildcat* (*Wildcat*LysA; UniProt entry Q19Y11)

**Figure 6.**
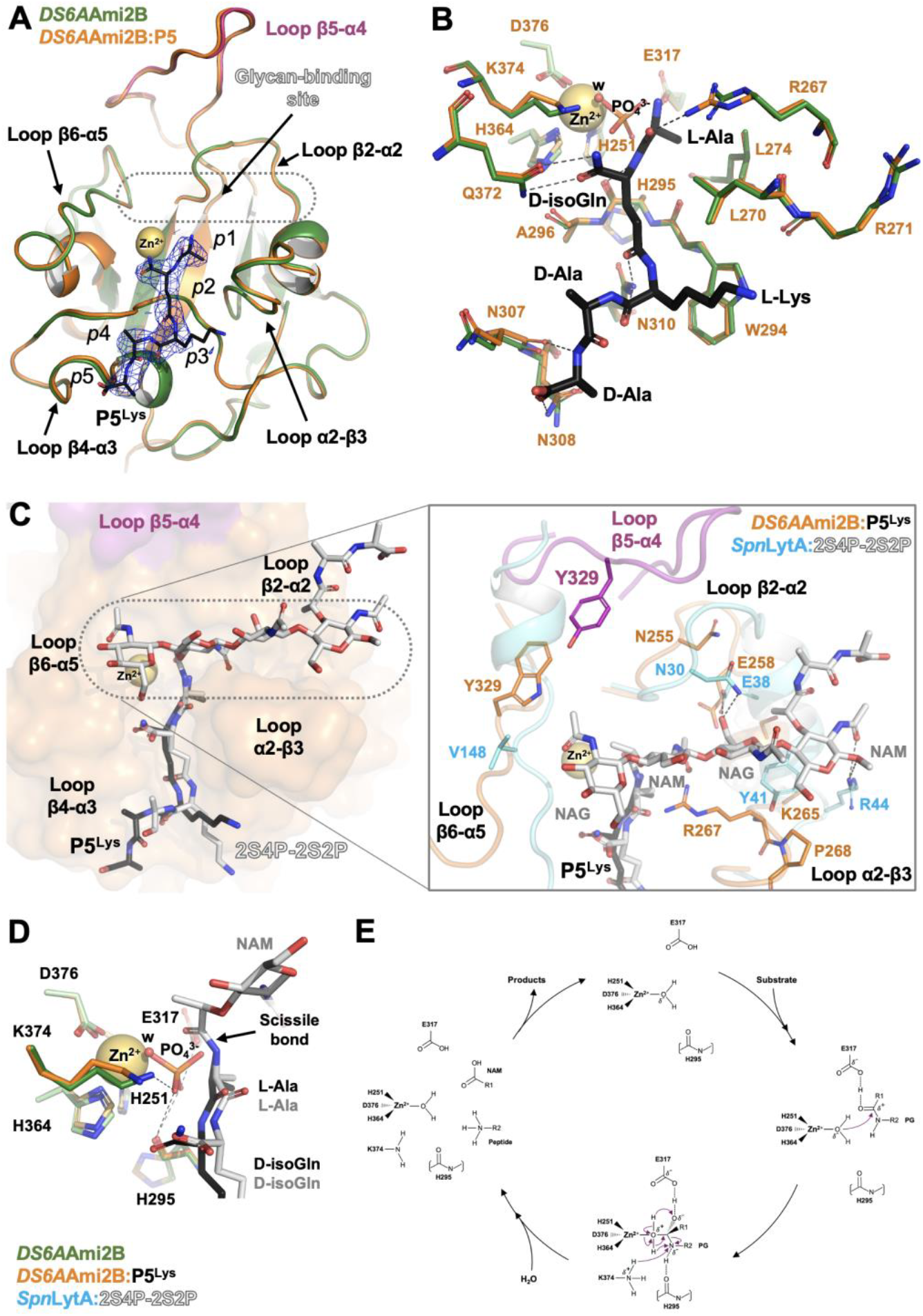
(A) Cartoon representation of the *DS6A*Ami2B:P5^Lys^ complex (in orange) superimposed to the native *DS6A*Ami2B structure (in green). Detail of the 2Fo-Fc electron density map (blue mesh) contoured at sigma=1.0 and defining the P5 ^Lys^ molecule (black sticks) is shown. The zinc cation bound in both structures is represented as a yellow sphere. The loops forming the both glycan and peptide-binding channels and *p*1-*p*5 amino-acid binding pockets are labelled. (B) View of the residues participating in the interaction with the P5^Lys^ molecule in *DS6A*Ami2B:P5^Lys^ and native *DS6A*Ami2B structures, coloured in orange and green, respectively. P5^Lys^ and Zn(II) are represented as in (A). (C) Superimposition of *DS6A*Ami2B:P5^Lys^ (in orange) and *Spn*LytA:4S5P (in cyan, PDB entry 5CTV) with P5^Lys^ and 4S5P molecules represented as black and white sticks, respectively. (D) Detail of the active sites in *DS6A*Ami2B bound to phosphate molecule, *DS6A*Ami2B:P5^Lys^ zinc-interacting water molecule, and *Spn*LytA:4S5P. (E) Proposed catalytic cycle in *DS6A*Ami2B.

#### 3.3.2. *DS6A*Ami2B substrate-binding can accommodate a tetrasaccharide-pentapeptide substrate

To investigate the molecular bases of substrate binding and hydrolysis, we determined the crystal structure of *DS6A*Ami2B in complex with the peptide L-Ala-D-*iso*-Gln-L-Lys-D-Ala-D-Ala (P5^Lys^ peptide), bearing a L-Lys residue instead of the mDAP found in the mycobacterial PG. The structure of *DS6A*Ami2B:P5^Lys^ was solved at 2.74 Å resolution, and it consists of three molecules of complex in the asymmetric unit exhibiting the same fold (RMSD of 0.139-0.176 Å for 165 Cα atoms), also conserved with respect to the native structure (RMSD of 0.190-0.230 Å for 175 Cα atoms) (**Fig. 6** and **6SM**). The three molecules of P5^Lys^ exhibit a conserved pose. The latter is unambiguously defined by the electron density map, but for the Cδ, Cε and Nξ side-chain atoms of L-Lys and the C-terminal carboxylate group exhibiting none or poor density (**Fig. 6AB**). The *DS6A*Ami2B: P5^Lys^ structure reveals a peptide-binding groove shaped by loops α2-β3, β4-α3 and β6-α5 and up to five pockets, from *p*1 to *p*5, accommodating each amino acid of the ligand (**Fig. 6AB**). The *p*1 pocket is formed by two leucine (L270 and L274) side-chains and R265, which stabilizing the methyl group and the main chain carboxyl group of the L-Ala. While L270 is conserved in all LysAs inspected, L274 and R265 are frequently substituted by Ala. In the latter case, a His residue located downstream (residue aa_i+2_) might play the role of R265. The *p*2 pocked interacts with the D-*iso*-Gln through an H-bond network involving W294 and H295 main-chains, and N310 and Q372 side-chains, with these latter exhibiting high conservation across the LysAs analysed (**Fig. 5C**). The mycobacterial PG may contain a D-*iso*-Glu or a D-*iso*-Gln in the second amino acid of the peptide moiety, which exhibit amide or carboxylate groups at the Cα atom of the residue, respectively (Vollmer *et al*, 2008). Interestingly, while Q372 forms a bidentate H-bond contact with the amide group of the D-*iso*-Gln, a carboxylate group would only enable a single H-bond interaction. However, and in such a case, this lost contact might be replaced by another H-bond interaction with K374, whose side chain locates near enough to the D-*iso*-Gln in our structure (**Fig. 6M**). Although further studies will be required to validate this proposal, our analysis points at *DS6A*Ami2B and LysA-Ami2B homologues as amidase domains able to recognize D-*iso*-Gln and D-*iso*-Glu at *p*2 of the peptide moiety. At *p*3, the side and main chains of L-Lys are stabilized by hydrophobic contacts formed with L270 and W294, and H-bond interactions with N310, respectively, all conserved across LysA-Ami2B domains. The side chain of L-Lys does not stablish specific contacts with Ami2B, which aligns with the flexibility observed for atoms Cδ, Cε and N (**Fig. 6**). This is not surprising, as the mycobacterial PG presents mDAP at the third amino-acid of the peptide moiety, which bears a carboxylate group absent in the L-Lys side-chain. Interestingly, inspection of residues nearby *p*3 does reveal the presence of an arginine (R271) adjacent to *p*3 as possible interactor of the mDAP side chain. At the end of the peptide-binding groove, the channel widens at *p*4 and *p*5 pockets that accommodate the D-Ala-D-Ala moiety via H-bond contacts of the ligand backbone with the N307 main-chain and the highly conserved N308 side-chain. Comparison of *DS6S*Ami2B:P5^Lys^ complex with the native structure does not show significant changes in the residues involved in P5^Lys^ interaction, thus suggesting the peptide-binding to be pre-formed for PG recognition. Nevertheless, we must bear in mind the carboxylate group missing in our peptide, due to the presence of L-Lys, which might affect potential rearrangements at loop α2-β3 driven by the proposed interactions of R271 with the mDAP side chain.

We also attempted the resolution of a *DS6A*Ami2B crystal structure bound to saccharide fragments. This did not yield any structure, which is likely due to steric hindrance of the glycan-binding site resulting of contacts with another molecule of the crystal. Thus, and to analyse the glycan-binding groove, we superimposed the *DS6A*Ami2B:P5^Lys^ complex with the crystal structure of the amidase domain of the *Spn*LytA autolysin in complex with a PG fragment (*Spn*LytA:4S5P, PDB entry 5CTV) (Sandalova *et al*, 2016) (**Fig. 6C**). This evidences a large glycan-binding channel in *DS6A*Ami2B able to accommodate up a tetra-saccharide NAG-NAM-NAG-NAM occupying positions −1, 0, +1 and +2. This between both glycan-binding channels. Related to the former, *DS6A*Ami2B presents a tryptophan residue (W329) edging the channel at −1 position in (V148 in *Spn*LytA), thus locking the substrate-binding channel beyond the −1 position. The W329 conformation is stabilised through contacts established with residues of a hydrophobic pocket, and through cation-pi contacts the arginine of an adjacent *DS6A*Ami2B:P5^Lys^ complex that is comparison also reveals significant differences and similitudes with additionally forming a salt-bridge interaction with a glutamate (E331) of the β5-α4 loop. The location and contacts stabilising W329 side-chain opens an interrogation regarding its conformation and potential flexibility in the isolated monomer. This and the conservation observed for W329 across LysAs suggest this residue as plausible element regulating the opening of the PG-binding channel, which would enable interactions with the glycan moiety beyond position −1 in LysA-Ami2B domains. This is an interesting posit that will require more studies to be validated. Other differences observed between *DS6A*Ami2B:P5^Lys^ and *Spn*LytA:4S5P complexes regards the α2 helix and the α2-β3 loop framing positions 0, +1 and +2. On the one hand, we observed narrower pockets in *DS6A*Ami2B at positions 0 and +1, where R267 would orient towards the NAM ring at position 0 to likely form favourable contacts. The R267 orientation would be safeguarded by restrictions imposed by a preceding proline and contacts stablished with the peptide moiety. On the other hand, a tyrosine residue found in other Amidase_2 members at position +2 (Y41 in *Spn*LytA) is missing in *DS6A*Ami2B and other LysA-Ami2B domains, thus resulting in a shallower +2 site. This seems to compensate the presence of a bulkier α2-β3 loop, thus resulting in enough room to accommodate the NAM and nascent peptide stem expected to bind at this position. Despite the differences observed, we identify three residues of *Spn*LytA (N30, E38 and R44) forming key contacts with the PG, and mutation of two of these residues has been shown to significantly decrease the catalytic activity of the enzyme (Table 4SM) (Mellroth *et al*, 2014). These residues are conserved across Amidase_2 domains, including *DS6A*Ami2B (N255, E258 and K265) and LysA-Ami2B inspected, thus pointing at a relevant role in PG binding by LysA-Ami2B amidases (**Fig. 5C**). Finally, the β5-α4 loop of *DS6A*Ami2B presents an aromatic residue (Y329), occupying a central position of the loop, that orients toward position −1. This residue is also found in other LysA-Ami2B domains, as tyrosine, phenylalanine and histidine (**Fig. 5C**). And in Amidase_2 homologues exhibiting a long β5-α4 loop such as *Srs*AmpDh2, where it has been shown to participate in substrate interaction at positions −1 and −2 (Martínez-Caballero *et al*, 2013), thus supporting the proposed PG-binding role of β5-α4 loop in LysA-Ami2B amidases (**Fig. 5C** and **6SM**).

To gain insights into *DS6A*Ami2B catalytic mechanism, we inspected the active site, in both native and peptide-bound structures, and compared it with the high-resolution structure of *Spn*LytA:S4P5 (**Fig. 6B**). This highlights, on the one hand, the zinc-interacting phosphate and water molecules laying adjacent the scissile bond in *DS6A*Ami2B native and complex structures, respectively. On the other hand, the phosphate lays adjacent the scissile bond in *DS6A*Ami2B and *Spn*LytA:S4P5 complexes mimicking the tetrahedral intermediate (TI) of the reaction to be formed upon the attack of the catalytic water (w^cat^) onto the lactyl group of the PG. In this line, the water molecule adjacent the scissile bond (< 3 Å) nicely overlays with one of the oxygens of the phosphate, thus emerging as the potential w^cat^ involved in catalysis. The oxyanion formed upon the catalytic attack is nicely mimicked by a second oxygen of the phosphate that locates at H-bond distance from the catalytic E317 side-chain, which supports E317 to function as an acid stabilising the TI. The TI formation would drag the peptide moiety towards the catalytic zinc, thus approaching the scissile bond towards His295 main-chain to a distance favourable for H-bond interactions, thus contributing to stabilise the TI state. In this configuration, the side chain of K374 would lay close enough to supply the proton necessary to form the amine group of the peptide product, thus favouring the final rupture of the substrate. While this catalytic mechanism partially aligns with the proposed for members of the Amidase_2 family (Zoll *et al*, 2010; Mellroth *et al*, 2014; Sandalova *et al*, 2016) it also features new posits regarding the roles played by E317 and K374 catalytic residues and assignment of the catalytic water in LysA-Ami2B and other Amidase_2 domains. Hypothesis that should be further investigated in order to unambiguously validate this mechanism.

### 3.4. Model of interaction of *DS6A*LysA catalytic domains with the mycobacterial cell-wall

We have determined the molecular envelop corresponding to the two catalytic domains of *DS6A*LysA in solution by SAXS and using the 6His-tagged and untagged versions of the *DS6A*GH19-Ami2B construct. Initially, we docked both crystallographic *DS6A*GH19 and *DS6A*Ami2B domains onto the molecular envelop calculated for the 6His-*DS6A*GH19-Ami2B construct. Using the protrusion resulting of the presence of the 6His-tag, we could unambiguously assign and dock each domain, which were oriented based on the location of the His-tag and the linker connecting GH19 and Ami2B domains (**Fig. 7SM**). The scattering curve calculated for this initial model shows a good fitting with the experimental data (**Fig. 7SM**). Thereafter, this initial model was docked onto the molecular envelop determined for the untagged *DS6A*GH19-Ami2B construct. *DS6A*GH19 and *DS6A*Ami2B domains were slightly readjusted in order optimise their fitting to the SAXS envelop, resulting in a model showing a good fitting with the experimental data (**Fig. 7** and **7SM**). Interestingly, examination of this final model shows that the PG-binding channels of both GH19 and Ami2B are facing the same side envelop, thus suggesting that both catalytic domains may orient towards a narrow region of the substrate to, concomitantly, proceed to its digestion (**Fig. 7**). To explore this hypothesis, we first superimposed the *D29*GH19^E228Q^:NAG_4_ and *DS6A*Ami2B:P5^Lys^ complexes onto the SAXS model. Next, we measured the distance between both glycan strands bound at *D29*GH19 and *DS6A*Ami2B PG-binding channels, which consisted of 40 Å approximately. Interestingly, the distance spanning glycan strands measured in a model of the PG molecule reported by NMR (Meroueh *et al*, 2006) falls very close to the one observed in our model, which suggest that both catalytic domains of *DS6A*LysA are likely to act sequentially in the hydrolysis of the PG molecule *in vivo* (**Fig. 7**).

**Figure 7.**
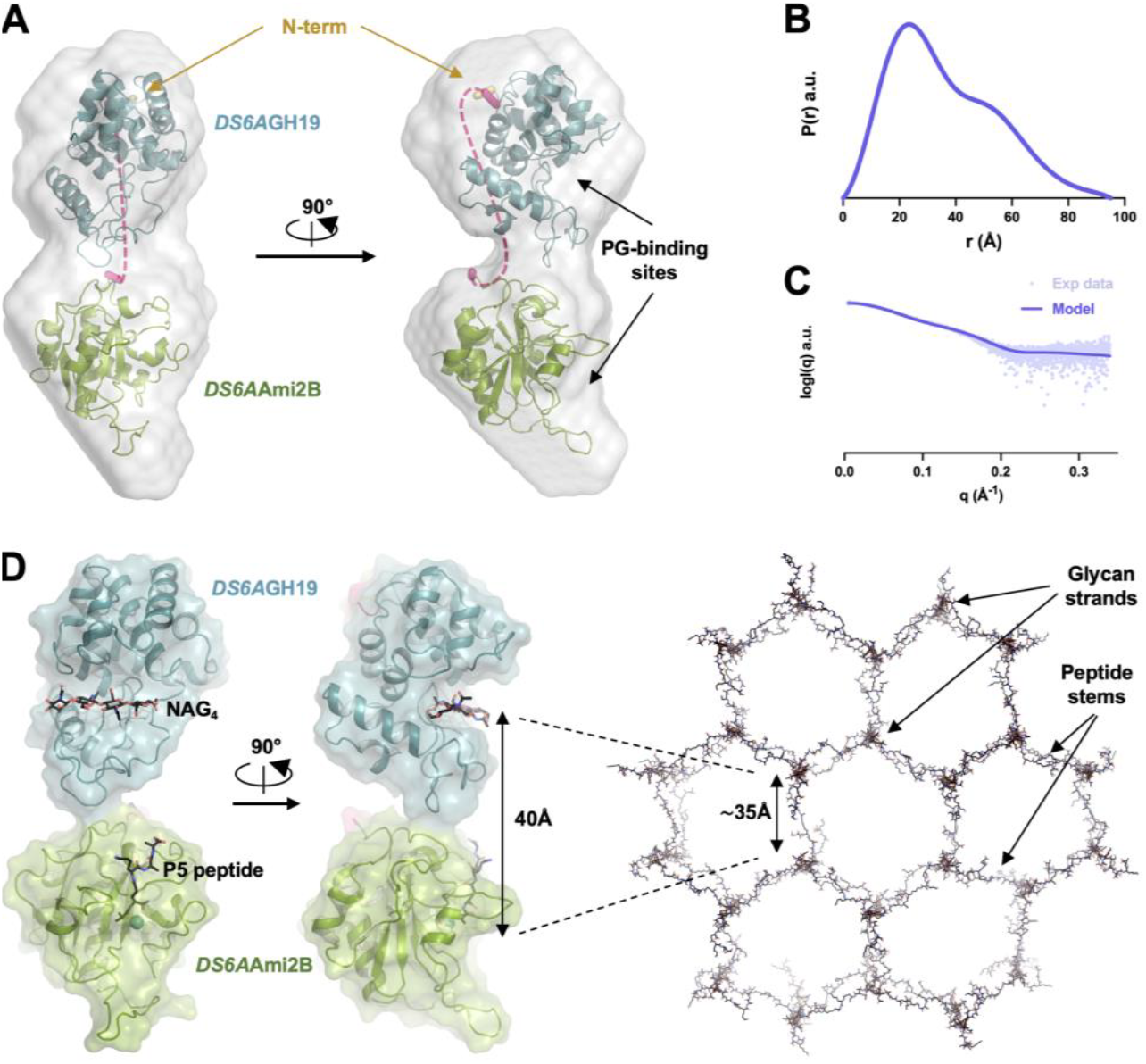
(A) *Ab initio* determined SAXS envelop for the untagged *DS6A*GH19-Ami2B construct in solution by SAXS (in grey) where the domains GH19 (in blue) and Ami2B (in green) have been fitted. The N-terminus of the GH19 domain (in yellow) is highlighted as thicker ribbons. The C-terminus of the GH19 domain and the N-terminus of the Ami2B domain (coloured in magenta) are highlighted as thicker ribbons. The dashed lines in pink do presume the location of the inter-domain linker. (B) Plot showing the normalized pair-distance distribution function P(r) for *DS6A*GH19-Ami2B (a.u., arbitrary units). (C) Experimental scattering curve (dots) and theoretical scattering curve computed for the GH19-Ami2B model (smooth). (D) View of the model corresponding of DS6ALysA catalytic domains, resulting of the superimposition of D29GH19:NAG4 (in blue) and *DS6A*Ami2B:P5^Lys^ (in green) complexes onto the SAXS model. Both PG fragments are depicted as black sticks, with the distance between their glycan moieties is indicated and compared with the dimensions of a model corresponding to the Gram-negative PG molecule determined by NMR (on the right).

## 4. Conclusions

Our work gathers fundamental details of peptidoglycan hydrolysis by *D29*LysA and *DS6A*LysA mycobacteriophage endolysins with therapeutic relevance for enzybiotics development to combat mycobacterial infections. Starting with the crystallographic characterisation of *D29*N4, our study has enabled us to experimentally validate the dipeptidyl-peptidase function of this domain and narrow its membership to the C41 family of peptidases, which aligns with previously reported *in silico* and biochemical studies. Importantly, we provide with new insights into the mechanism of PG binding and hydrolysis by *D29*N4, including the precise arrangement of the active site and the identification of the β1-β2 loop as a potential regulatory element in peptidoglycan interaction. Moreover, and based on biochemical studies reported elsewhere, we have mapped key residues for substrate interaction. Also, we have identified new residues potentially involved in the stabilisation of the mDAP moiety of substrate, as a particular feature of this domain with implications in substrate selectivity. We also show conservation of these structural-functional features in other homologous LysA proteins, thus shedding light into the function of this class of N4 peptidases.

Our study of *D29*GH19 and *DS6A*GH19 shows that LysA-GH19s domains are “loopless” glycosyl-hydrolases belonging to the GH19_1 subfamily, and characterised by smaller substrate-binding channels when compared to their “loopful” chitinase homologues belonging to the GH19_2 subfamily. Our crystallographic characterisation of several *D29*GH19 complexes with glycan analogues has enabled us to capture snapshots of the catalytic process. The latter transits from an initial open state of the domain (*D29*GH19^E228Q^:NAG_4_), necessary for substrate binding but not suitable for hydrolysis, to a closed state of the domain (*D29*GH19^WT^:NAG:NAG_2_ and *D29*GH19^WT^:NAG) indispensable for correct substrate binding and hydrolysis, which we have captured as post-hydrolysis stages *in crystallo*. Our structural analysis evidences the significant inter-lobe rearrangement required for this transition that only occurs when the enzyme is catalytically ready. Moreover, our data supports the idea that the M-lobe functions as a pre-formed structural platform for substrate interaction in the open state, positioning the substrate correctly towards its catalytic base. This crucial step is followed by the closure of the PG-binding groove around the substrate to enable a correct approximation of the catalytic acid towards the glycosidic bond, in line with the GH19 family hydrolase mechanism with inversion of the configuration (Iseli et al, 1996). Additionally, a detailed analysis of our crystallographic and docking structures has led us to identify essential residues involved in PG binding and hydrolysis, as well as evidence supporting *D29*GH19 to operate as muramidase enzyme cleaving the glycosidic bond connecting NAM and NAG saccharides of the PG. These features extend to other LysA-GH19 domains, thus unravelling key characteristics for this class of endolysin domains and for the whole subfamily of GH19_1 hydrolases.

We also report the crystal structures of the *DS6A*Ami2B domain, alone and in complex with an analogue of the product of the reaction. Our structural analysis evidence *DS6A*Ami2B as member of the Amidase_2 family, conserving the characteristic catalytic features described for these group of amidases. Our structures reveal central details of the mode of interaction of this enzyme with the peptide moiety of the PG, which occurs through a large cavity capable to accommodate a substrate of up to four saccharide rings and five amino acids of the peptide stem. We also uncover particular features of the PG-binding channel of *DS6A*Ami2B in comparison to other Amidase_2 homologues, such as: i) a rigid loop α2-β3 loop preforming *p*1, *p*2, 0 and +1 pockets; ii) the presence of a tryptophan locking the potion −2 of the glycan-binding groove, which emerges as a potential gatekeeper regulating accessibility of the PG-biding site; and iii) the presence of a long β5-α4 loop, potentially involved in interaction with the glycan moiety. Moreover, a phosphate molecule mimicking the tetrahedral intermediate state of the reaction in the native structure has allowed us to explain potential interactions at the active site during catalysis, enabling us to propose a model of the catalytic mechanism for *DS6A*Ami2B and other LysA-Ami2B domains. In this study, we have further explored the arrangement of *DS6A*LysA catalytic domains in solution using SAXS. In order to full-proof the relative orientation of both muramidase and amidase domains, we characterised the enzyme in-solution by SAXS with and without the presence of the His-tag. The results permitted us to understand the manner and orientation adopted by both domains with respect to each other. Our SAXS model supports both PG-binding sites to be located at the same side of the molecular envelop, thus supporting both catalytic activities functioning together and explaining a sequential model of digestion of the cell-wall. Altogether, the present study constitutes a solid groundwork for future investigations on this relevant class of endolysins, including the rational design of LysA lysins with optimised enzybiotic activity to treat mycobacterial IDs and to combat emergent antimicrobial resistance associated to this class of pathogens.

## Acknowledges

This work was supported by a Atracción de Talento program (Modalidad 1) grant from the Madrid Regional Government (Spain). The authors would like to also thank Prof. Martín Martínez-Ripoll and Prof. Juan A. Hermoso for their precious discussion about this work and manuscript revision; to Prof. Shahriar Mobashery for kindly sharing with us the coordinate file corresponding to the PG structural-model; and to Dr. Beatriz González as MB-BAG coordinator, for facilitating us with access to MX beamtime at the European Synchrotron Radiation Facility (ESRF). We would like to thank the ESRF (beamline ID30A-3, proposals mx33176 and mx35926), Diamond Light Source (beamline B21, proposals mx33176 and mx35926) and the ALBA Synchrotron (XALOC-BL13 beamline proposals 2021085244 and 2022097082) for beamtime and their staff for assistance during data collection.

## Author contribution

I.P.D. conceived the project and supervised the experiments. F.C.Z. performed the expression, purification, structural characterization of (PDB entries of structures 9HNV, 9HNU, 9HP7, 9HNA and 9HTY). L.G.L. performed the expression, purification, structure characterization of (PDB entries of structures 9HYR, 9HQW, 9HR1, 9HRM, 9HV2 and 9HV0). F.C.Z. carried out the final validation and structure deposition of all the crystallographic structures in the Protein Data Bank. I.P.D. carried out the analysis of the crystallographic structures with the contribution of F.C.Z. and L.G.L.

I.G.M. performed the SAXS studies, including data analysis, figure preparation and data deposition. L.I.S. carried out the computational docking studies and model analysis. I.P.D. wrote the manuscript and prepared the figures with involvement of all the other authors.

## Supplementary Material Figures

**Figure 2SM.**
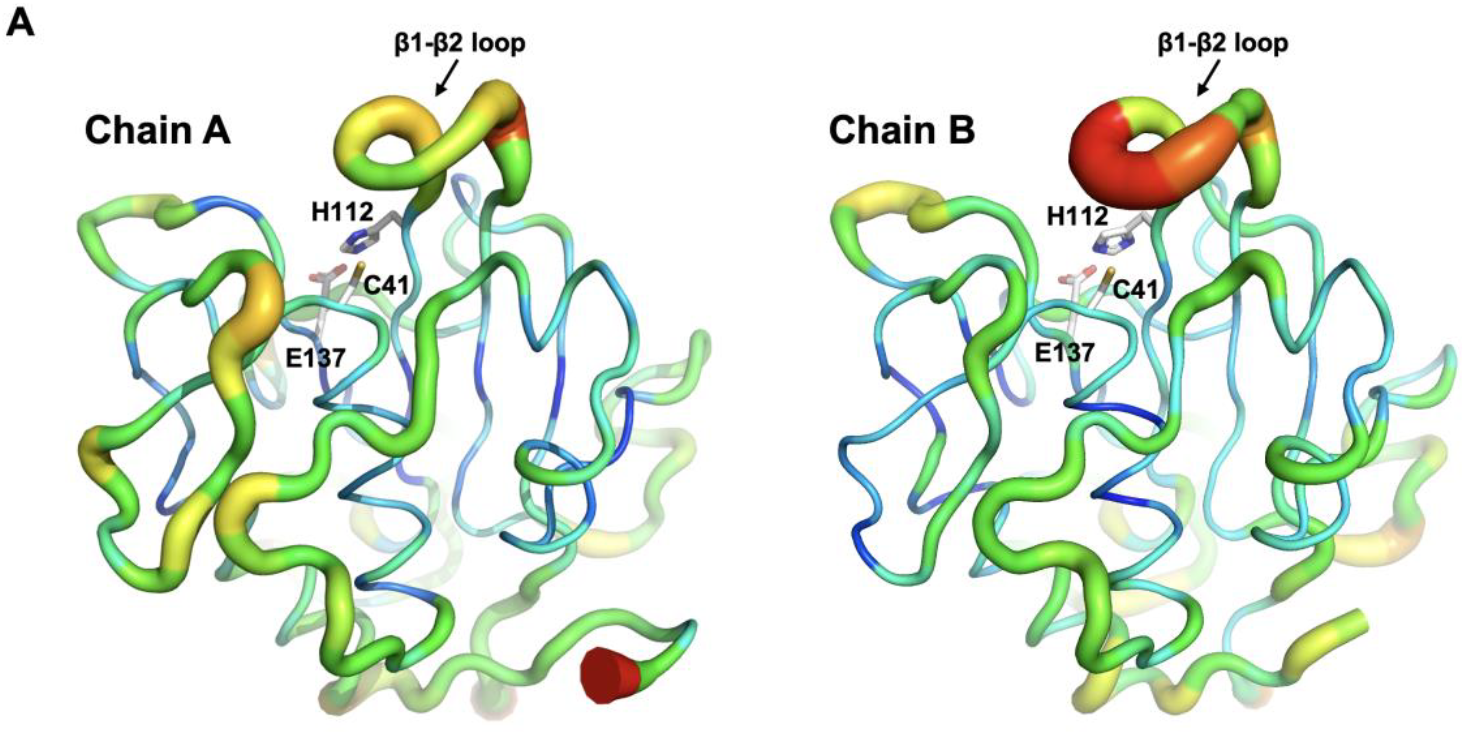
Normalised representation of the B-factor values observed in chains A (left) and B (right) of the crystal structure of *D29*N4, which range from colour blue (lowest B-factor values) to red (highest B-factor values). The catalytic triad (represented as sticks) and the β1-β2 loop are indicated with a labels.

**Figure 3SM.**
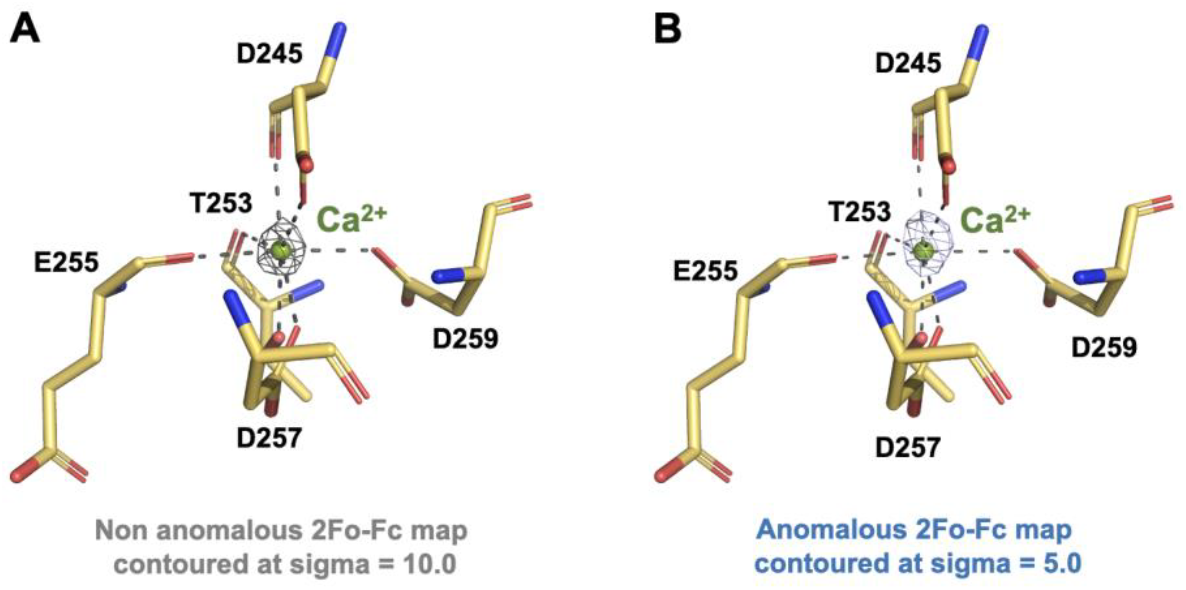
Detail of the EF-hand-like calcium-binding site of *D29*GH19. Electron density peaks corresponding to the calcium cation identified at both non anomalous and anomalous 2Fo-Fc maps are depicted as a mesh.

**Figure 4SM.**
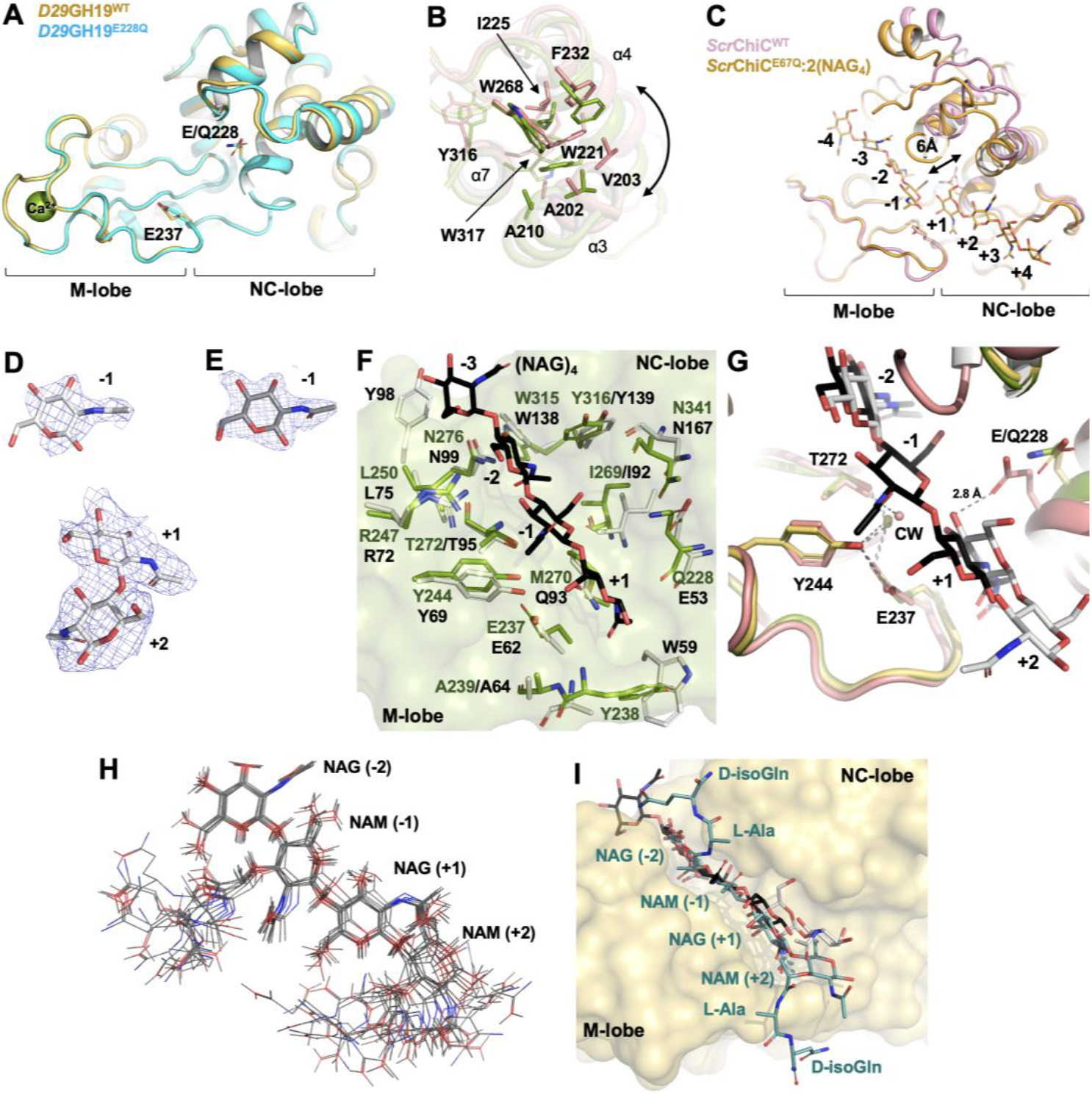
(A) Superimposition of *D29*GH19^WT^ and *D29*GH19^E288Q^ represented as cartoons in yellow and cyan, respectively, depicting the catalytic residues as sticks. (B) Detail of the hydrophobic core involved in NC-lobe pivoting towards α7 between *D29*GH19^E228Q^:NAG_4_ (in green) and *D29*GH19^WT^:NAG:NAG_2_ (in pink). (C) Superimposition the of crystal structures of *Rsc*-C, in the absence (PDB entry 4DWX) and bound to a NAG_4_ (PDB entry 4J0L), illustrating the closure of the substrate-binding cavity upon ligand binding in a “loop-full” GH19 domain. (D) and (E) detail of the 2Fo-Fc electron density map (blue mesh) corresponding to the NAG:NAG_2_ and NAG molecules bound in complexes *D29*GH19^WT^:NAG (white) and *D29*GH19^WT^:NAG:NAG_2_ (grey), respectively, countered at sigma = 1.0. (F) Superimposition of *D29*GH19^E228Q^:NAG_4_ (protein in green, NAG_4_ in black) and *DS6A*GH19 (white) structures showing residues involved in substrate interaction in both domains as sticks. (G) Detail of the active site in *D29*GH19^WT-I^, *D29*GH19^E228Q^:NAG_4_ and *D29*GH19^WT^:NAG:NAG_2_ superimposed structures. Key residues involved in catalysis and NAG_n_ fragments are represented as sticks following the same colour code as in (A). The catalytic water (CW) molecule activated by E237 is represented as spheres coloured in yellow (in *D29*GH19^WT-I^), green (in *D29*GH19^E228Q^:NAG_4_) and pink (in *D29*GH19^WT^:NAG:NAG_2_). (H) Representation of the eleven top solutions obtained in the docking of 4S2P PG-fragment (lines coloured in grey) onto *D29*GH19^WT^. (I) Superimposition of the top docking solution calculated for *D29*GH19^WT^:4S2P complex (protein in yellow, ligand in blue) with ligands bound to the crystallographic complexes *D29*GH19^E228Q^:NAG_4_ (black sticks) and *D29*GH19^WT^:NAG:NAG_2_ (white sticks).

**Figure 6SM.**
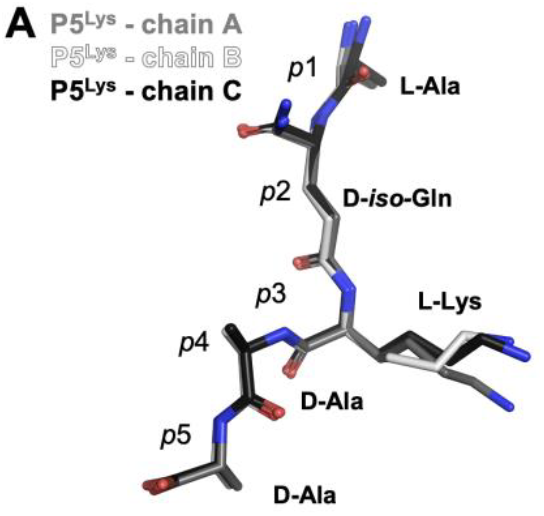
(A) Superimposition of the three complexes showing the peptide pose.

**Figure 7SM.**
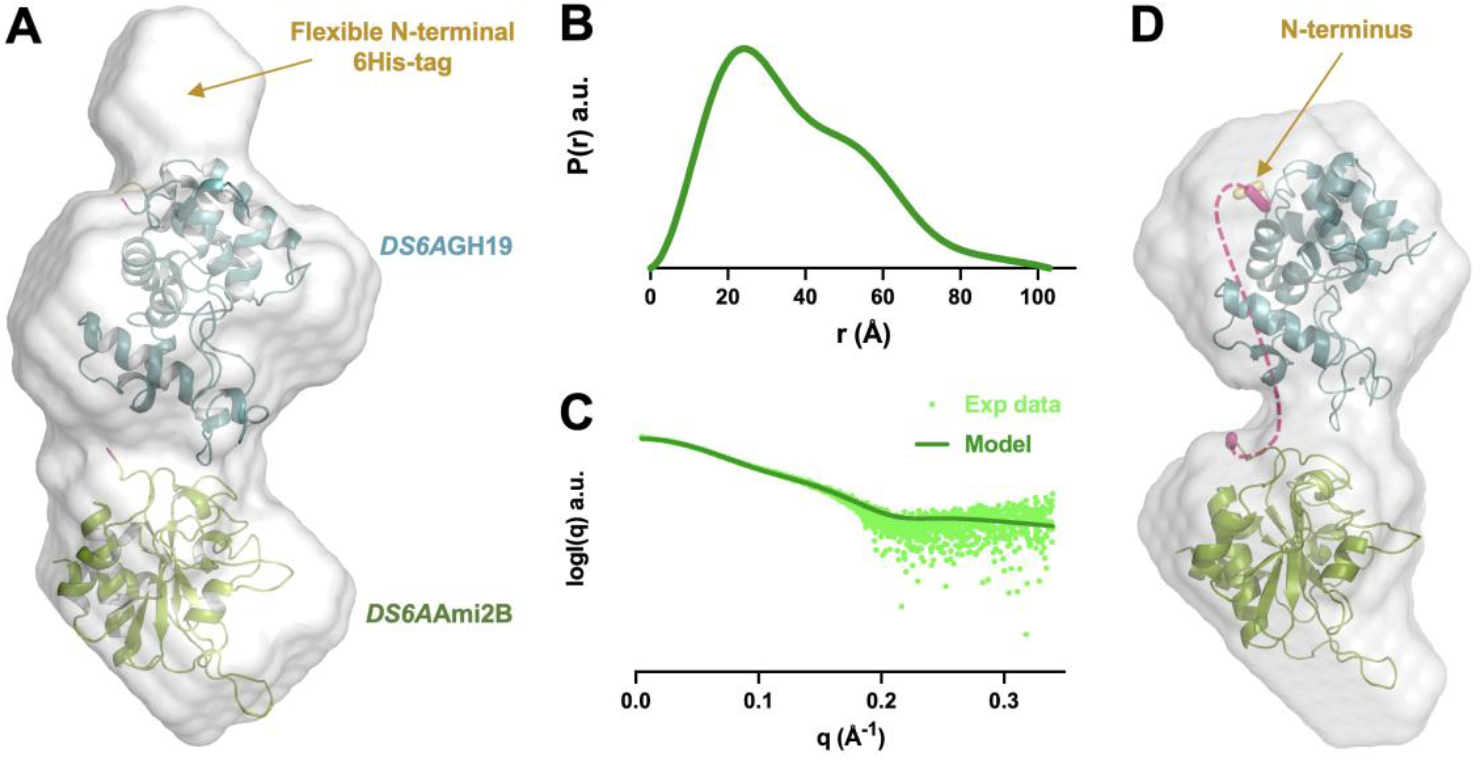
(A) *Ab initio* molecular envelope determined by SAXS for a *DS6A*Ami2B construct containing a N-terminal 6His-tag. (B) Normalized pair-distance distribution function P(r) for the protein without the his-tag (green graph). (C) Experimental scattering curve (lime green dots) and theoretical scattering curve, in emerald green, computed for the model (smooth), where the protein adopts a conformation that could be explained by the model resulting from joining the PDBs 9HQW and 9HNA, using the program Coot. The data shows an extra volume that clarifies where the N-terminal domain of *DS6A*GH19 is located with respect to the *DS6A*Ami2B domain. (D) *Ab initio* molecular envelope determined by SAXS for the untagged DS6AAmi2B construct.

**Table 1SM.** X-ray data collection and refinement statistics.

|  | *D29N4<sup>WT</sup>*<br>*native* | *D29GH19<sup>WT</sup>*<br>*native*<br>(Form I) | *D29GH19<sup>WT</sup>*<br>*native*<br>(Form II) | *D29GH19<sup>WT</sup>*<br>*native*<br>(Form III) | *D29GH19<sup>WT</sup>*<br>with NAG<sub>2</sub>/NAG<br>(Form IV) | *D29GH19<sup>WT</sup>*<br>with NAG<br>(Form III) | *DS6AGH19<sup>WT</sup>*<br>*native* | *D29GH19<sup>E228Q</sup>*<br>*native*<br>(Form V) | *D29GH19<sup>E228Q</sup>*<br>with NAG<sub>4</sub><br>(Form I) | *DS6AAmi2B<sup>WT</sup>*<br>*native* | *DS6AAmi2B<sup>WT</sup>*<br>with P5<sup>lys</sup> |
| --- | --- | --- | --- | --- | --- | --- | --- | --- | --- | --- | --- |
| Data collection statistics |  |  |  |  |  |  |  |  |  |  |  |
| PDB entry code | 9HYR | 9HQW | 9HR1 | 9HRM | 9HV2 | 9HV0 | 9HNV | 9HNU | 9HP7 | 9HNA | 9HTY |
| Space group | *P*2<sub>1</sub>2<sub>1</sub>2<sub>1</sub> | *C*2 | *P*2<sub>1</sub>2<sub>1</sub>2<sub>1</sub> | *P*2<sub>1</sub>2<sub>1</sub>2<sub>1</sub> | *P*2<sub>1</sub>2<sub>1</sub>2 | *P*2<sub>1</sub>2<sub>1</sub>2<sub>1</sub> | *I*222 | *I*2 | *C*2 | *C*2 | *C*2 |
| No. monomers in ASU | 2 | 1 | 2 | 2 | 4 | 2 | 3 | 1 | 1 | 3 | 3 |
| *a*, *b*, *c* (Å) | 65.2, 67.0, 83.0 | 91.3, 44.8, 69.7 | 44.6, 89.0, 113.4 | 37.0, 57.0, 151.3 | 78.7, 84.2, 135.3 | 37.2, 56.5, 152.6 | 89.2, 145.9, 147.7 | 70.1, 44.9, 76.6 | 91.1, 44.8, 69.5 | 72.0, 121.0, 80.7 | 70.7, 121.2, 80.0 |
| $\alpha$, $\beta$, $\gamma$ (°) | 90.0, 90.0, 90.0 | 90.0, 126.1, 90.0 | 90.0, 90.0, 90.0 | 90.0, 90.0, 90.0 | 90.0, 90.0, 90.0 | 90.0, 90.0, 90.0 | 90.0, 90.0, 90.0 | 90.0, 103.0, 90.0 | 90.0, 126.4, 90.0 | 90.0, 107.2, 90.0 | 90.0, 106.3, 90.0 |
| Wavelength (Å) | 0.9793 | 0.9793 | 0.9793 | 0.9677 | 0.9677 | 0.9677 | 0.9792 | 0.9792 | 0.9792 | 0.9793 | 0.9792 |
| Beamline | XALOC | XALOC | XALOC | ID30A-3 | ID30A-3 | ID30A-3 | XALOC | XALOC | XALOC | XALOC | XALOC |
| Resolution range (Å) | 82.97-3.20<br>(3.42-3.20) | 56.27-1.63<br>(1.66-1.63) | 113.39-2.00<br>(2.05-2.00) | 57.04-2.02<br>(2.10-2.04) | 45.81-3.10<br>(3.31-3.10) | 76.31-2.43<br>(2.52-2.43) | 76.34-2.75<br>(2.88-2.75) | 45.56-1.45<br>(1.47-1.45) | 38.25-1.83<br>(1.86-1.83) | 77.14-2.04<br>(2.07-2.04) | 60.62-2.74<br>(2.79-2.74) |
| *R*<sub>merge</sub> | 0.321 (0.701) | 0.059 (0.597) | 0.117 (0.895) | 0.220 (1.041) | 0.271 (1.448) | 0.174 (0.891) | 0.344 (2.156) | 0.047 (0.312) | 0.073 (0.758) | 0.134 (1.240) | 0.112 (0.415) |
| *R*<sub>pim</sub> | 0.173 (0.376) | 0.036 (0.388) | 0.033 (0.246) | 0.160 (0.754) | 0.136 (0.710) | 0.097 (0.486) | 0.128 (0.799) | 0.021 (0.158) | 0.031 (0.346) | 0.055 (0.497) | 0.072 (0.258) |
| No. total reflections | 27415 (4952) | 101079 (4353) | 434589 (33922) | 81875 (6595) | 73573 (13776) | 47708 (5232) | 210189 (27695) | 236632 (9170) | 126584 (5698) | 283455 (14979) | 56465 (2970) |
| No. unique reflections | 6267 (1110) | 27799 (1355) | 30434 (2255) | 20836 (1595) | 16002 (2889) | 12304 (1286) | 25458 (3356) | 41013 (1973) | 19241 (991) | 41716 (2098) | 16754 (845) |
| Mean (*I*)/SD(*I*) | 4.7 (2.4) | 14.2 (2.2) | 17.2 (3.5) | 5.5 (1.8) | 5.2 (1.2) | 6.1 (1.8) | 6.7 (1.2) | 16.8 (3.2) | 15.1 (2.2) | 10.2 (2.1) | 5.6 (2.3) |
| CC<sub>1/2</sub> | 0.934 (0.728) | 0.998 (0.816) | 0.999 (0.880) | 0.979 (0.514) | 0.985 (0.523) | 0.983 (0.516) | 0.984 (0.512) | 0.999 (0.970) | 0.999 (0.872) | 0.994 (0.666) | 0.991 (0.864) |
| Completeness (%) | 99.1 (99.7) | 98.4 (97.6) | 96.9 (100.0) | 98.5 (99.6) | 95.2 (96.4) | 96.6 (97.9) | 100.0 (100.0) | 99.3 (97.6) | 95.1 (99.9) | 98.7 (97.9) | 98.0 (99.8) |
| Multiplicity | 4.4 (4.5) | 3.6 (3.2) | 14.3 (15.0) | 3.9 (4.1) | 4.6 (4.8) | 3.9 (4.1) | 8.3 (8.3) | 5.8 (4.6) | 6.6 (5.7) | 6.8 (7.1) | 3.4 (3.5) |
| Refinement statistics |  |  |  |  |  |  |  |  |  |  |  |
| Resolution range (Å) | 51.25-3.20 | 45.49-1.64 | 70.04-2.00 | 53.37-2.04 | 43.81-3.10 | 76.31-2.43 | 76.34-2.75 | 45.56-1.45 | 38.25-1.83 | 77.14-2.04 | 60.62-2.74 |
| No. reflections |  | 27776 | 30368 | 20789 | 15962 | 12264 | 25438 | 40948 | 19225 | 41701 | 16749 |
| *R*<sub>work</sub> (%) | 0.25 | 0.18 | 0.17 | 0.18 | 0.24 | 0.21 | 0.24 | 0.15 | 0.18 | 0.18 | 0.16 |
| *R*<sub>free</sub> (%) | 0.31 | 0.21 | 0.20 | 0.24 | 0.28 | 0.26 | 0.28 | 0.18 | 0.20 | 0.21 | 0.20 |
| No. of atoms | 2714 | 1763 | 3463 | 2922 | 6195 | 2740 | 4630 | 1770 | 1688 | 4663 | 4515 |
| Protein | 2714 | 1515 | 3064 | 2681 | 6007 | 2682 | 4535 | 1542 | 1519 | 4368 | 4326 |
| Ion | — | 1 | 2 | — | 4 | — | — | 1 | 1 | 28 | 18 |
| PG analogue | — | — | — | — | 180 | 15 | — | — | 57 | — | 102 |
| Water | — | 227 | 355 | 241 | 8 | 43 | 95 | 213 | 111 | 267 | 69 |
| Other (MES, PEG, MPD, Ala, D-Ser) | — | 12/_/8/_ | 24/12/_/6 | — | — | — | — | 14 | — | — | — |
| B factors (Å<sup>2</sup>) |  |  |  |  |  |  |  |  |  |  |  |
| Protein | 34.5 | 23.9 | 28.4 | 21.5 | 62.0 | 32.2 | 44.0 | 24.4 | 30.87 | 36.81 | 34.35 |
| Other (MES, PEG, MPD, Ala, D-Ser) | — | 21.8/_/34.5/_ | 34.4/38.1/_/35.1 | — | — | — | — | 46.6 | — | — | — |
| PG analogue | — | — | — | — | 64.1 | 29.0 | — | — | 43.86 | — | 43.43 |
| Ion | — | 16.1 | 24.4 | — | 71.9 | — | — | 17.33 | 21.73 | 28.02 | 27.48 |
| Water | — | 34.4 | 36.9 | 29.2 | 59.6 | 32.2 | 36.41 | 34.78 | 36.68 | 43.78 | 34.96 |
| R.m.s. deviations |  |  |  |  |  |  |  |  |  |  |  |
| Bond lengths (Å) | 0.002 | 0.005 | 0.006 | 0.006 | 0.002 | 0.003 | 0.003 | 0.011 | 0.004 | 0.005 | 0.005 |
| Angle lengths (°) | 0.545 | 0.750 | 0.735 | 0.776 | 0.446 | 0.666 | 0.614 | 1.103 | 0.659 | 0.788 | 0.741 |

**Table 2SM.**
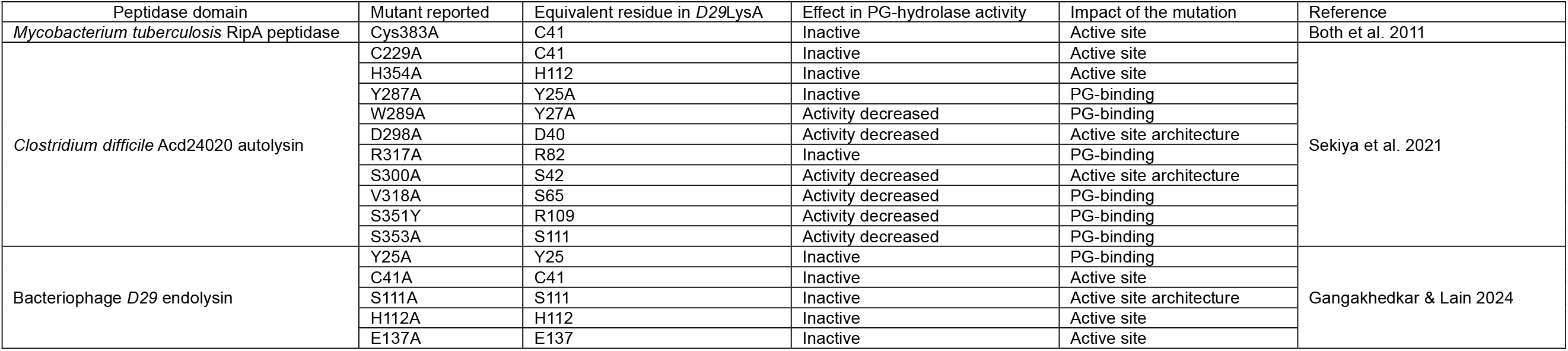
Mutants reported in *D29*N4 and homologue domains.

**Table 3SM.**
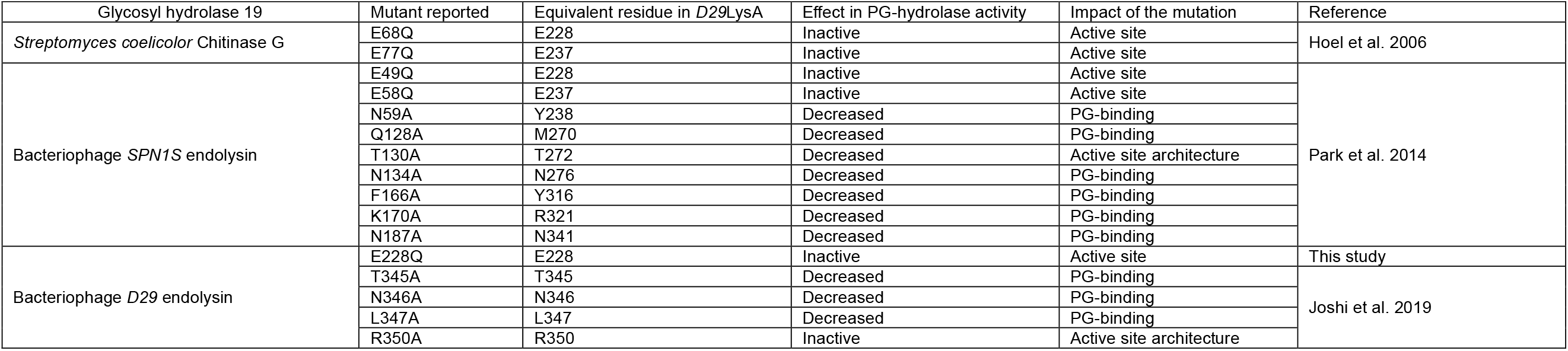
Mutants reported in *D29*GH19 and homologue domains.

**Table 4SM.**
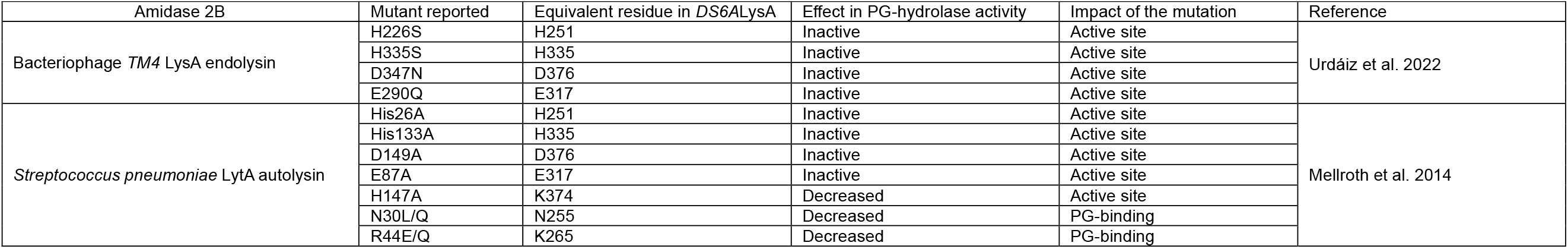
Mutants reported in *DS6A*Ami2B homologue domains.

| Amidase 2B | Mutant reported | Equivalent residue in DS6ALysA | Effect in PG-hydrolase activity | Impact of the mutation | Reference |
| --- | --- | --- | --- | --- | --- |
| Bacteriophage TM4 LysA endolysin | H226S | H251 | Inactive | Active site | Urdáiz et al. 2022 |
|  | H335S | H335 | Inactive | Active site |  |
|  | D347N | D376 | Inactive | Active site |  |
|  | E290Q | E317 | Inactive | Active site |  |
| *Streptococcus pneumoniae* LytA autolysin | His26A | H251 | Inactive | Active site | Mellroth et al. 2014 |
|  | His133A | H335 | Inactive | Active site |  |
|  | D149A | D376 | Inactive | Active site |  |
|  | E87A | E317 | Inactive | Active site |  |
|  | H147A | K374 | Decreased | Active site |  |
|  | N30L/Q | N255 | Decreased | PG-binding |  |
|  | R44E/Q | K265 | Decreased | PG-binding |  |

**Table 5SM.** SAS sample details, data collection, analysis, and 3D modelling details for biomolecules in solution.

|  |  |  |
| --- | --- | --- |
| Scattering particle composition |  |  |
| Protein | Untagged DS6AGH19-Ami2B | His-tagged DS6AGH19-Ami2B |
| Sample environment/configuration |  |  |
| Solvent composition | 20 mM Tris pH 8.0, 150 mM NaCl, 1 mM DTT |  |
| Sample temperature (°C) | 4 |  |
| In beam sample cell (°C) | 18 |  |
| Size Exclusion Chromatography SEC-SAS |  |  |
| Sample injection concentration, (mg/ml) | 5.9 | 5.6 |
| Sample injection volume, mL | 45 |  |
| SEC column type | SRT-C SEC-300 (Sepax) | Shodex KW-403 |
| SEC flowrate, mL/min | 0,16 | 0.16 |
| Data acquisition/reduction software | GDA/Scätter | GDA/Scätter |
| Source/instrument description or reference | B21, Diamond Light Source | B21, Diamond Light Source |
| Measured q-range ($q_{\min} - q_{\max}$; Å<sup>-1</sup>) | 0.0031-0.37 | 0.0031-0.37 |
| Exposure time (s) and final number of sample frames | 3, 600 | 3, 600 |
| Relevant details |  |  |
| Methods/Software | PRIMUS, AUTORG, GNOM |  |
| Guinier Analysis |  |  |
| I (0) ± σ (cm<sup>-1</sup>) | 0,03900 | 0,03400 |
| R<sub>g</sub> ± σ (Å) | 28,45±0,04 | 28,43±0,06 |
| Data point range | 80-323 | 80-315 |
| PDDF/P(r) analysis |  |  |
| I (0) ± σ (cm<sup>-1</sup>) | 0,03921 | 0,03397 |
| R<sub>g</sub> ± σ (Å) | 27,89±0,04 | 29,15±0,03 |
| d<sub>max</sub> (Å) | 95 | 103 |
| Data point range | 54-2164 | 28-1600 |
| q-range (Å<sup>-1</sup>) | 0,014-0,2866 | 0,0080-0,2132 |
| Methods/Software |  |  |
| Volume estimates |  |  |
| Porod volume, V<sub>p</sub> (Å<sup>3</sup>) | 500,900 | 621,000 |
| Molecular weight (M) estimates (kDa) |  |  |
| From chemical composition | 43.5 | 45,4 |
| From SAS, concentration independent method (MoW) | 39,8 | 49,2 |
| From SAS-independent measure (MALLS) | XXXX | XXXX |
| Shape modelling method |  |  |
| Software | DAMMIN, DAMMIF, FoXS |  |
| q-range for fit ($q_{\min} - q_{\max}$; Å<sup>-1</sup>) | 0,0045-0,3398 | |
| Symmetry/anisotropy assumptions | P1 | P1 |
| Number of individual model reconstructions | 20 | 20 |
| χ<sup>2</sup>, CorMap P-values for fit | 1,44, 1,03 | 0,64, 1.05 |
| Atomistic modelling methods |  |  |
| Software | FoXS |  |
| q-range for fit ($q_{\min} - q_{\max}$; Å<sup>-1</sup>) | 0,0045-0,3398 | 0,0045-0,3398 |
| Symmetry/anisotropy assumptions | N/A | N/A |
| Number of individual model reconstructions | 1 (Based on PDBs: 9HNA y 9HNV) | 1 (Based on PDBs: 9HNA y 9HNV) |
| χ<sup>2</sup>, CorMap P-values for fit | 0,90, 1,04 | 1,24, 1,02 |
| Data and model deposition |  |  |
| SASBDB IDs | SASDXXX ID 6560 | SASDXXX ID 6561 |

